# Florigen family chromatin recruitment, competition and target genes

**DOI:** 10.1101/2020.02.04.934026

**Authors:** Yang Zhu, Samantha Klasfeld, Cheol Woong Jeong, Run Jin, Koji Goto, Nobutoshi Yamaguchi, Doris Wagner

## Abstract

Plants monitor seasonal cues, such as day-length, to optimize life history traits including onset of reproduction and inflorescence architecture ^1–3^. Florigen family transcriptional co-regulators TERMINAL FLOWER 1 (TFL1) and FLOWERING LOCUS T (FT) antagonistically regulate these vital processes ^4–6^ yet how TFL1 and FT execute their roles and what the mechanism is for their antagonism remains poorly understood. We show genome-wide, that TFL1 is recruited to the chromatin by the bZIP transcription factor FLOWERING LOCUS D (FD) in *Arabidopsis*. We find that seasonal cue-mediated upregulation of FT competes TFL1 from chromatin-bound FD at key target loci. We identify the master regulator of floral fate, *LEAFY* (*LFY*) as a target under dual opposite regulation by TFL1 and FT. Exonic bZIP motifs in *LFY* are critical for repression by TFL1, upregulation by FT and adoption of floral fate. Transcriptomic identification of target genes directly repressed by the TFL1-FD complex not only identifies key regulators of onset of reproduction and floral fate, but reveals that TFL1-FD repress sugar and hormone signalling pathways and chromatin regulators. Our data provide mechanistic insight into how florigen family member sculpt inflorescence architecture, a trait important for reproductive success and yield.

## Main

Plant development occurs after embryogenesis and is plastic, this allows modulation of the final body plan in response to environmental cues to enhance growth and reproductive success ^7, 8^. Of particular importance for species survival is the timing of the formation of flowers that give rise to seeds which occurs in response to the seasonal cues photoperiod and temperature ^1, 3, 9^. For example, in plants that flower only once, like *Arabidopsis* and most crops, an early switch to flower formation allows rapid completion of the life-cycle in a short growing season, but reduces total seed set or yield ^10–12^. By contrast, delaying flower formation supports formation of more seeds, but extends the time to seed set. In many plant species, these alternative developmental trajectories are tuned in response to daylength in antagonistic fashion by two members of the florigen family of proteins ^12, 13^. FT promotes onset of the reproductive phase and flower formation (determinacy), while TFL1 promotes vegetative development and branch fate (indeterminacy) ^4–6, 13^. In *Arabidopsis*, which flowers in the spring, FT accumulates only when the daylength exceeds a critical threshold, while TFL1 is present in both short-day and long-day conditions ^1, 3, 14^. A key unanswered question is how FT and TFL1 modulate plant form – what are the downstream processes they set in motion and what is molecular basis for their antagonism?

Accumulating evidence points to roles of FT and TFL1, which lack ability to bind DNA, in transcriptional activation and repression, respectively, by forming complexes with a bZIP transcription factor, FLOWERING LOCUS D (FD) ^12, 15–19^, although non-nuclear functions for both proteins have also been described ^20, 21^. Mechanistic insight into florigen activity has been hampered by their low protein abundance. To overcome this limitation and to test the role of TFL1 in the nucleus, we conducted TFL1 chromatin immunoprecipitation followed by sequencing (ChIP-seq). Towards this end, we first generated a biologically active, genomic GFP-tagged version of TFL1 (gTFL1-GFP *tfl1-1*) (**Supplementary Fig. 1a, b**). Next, we enriched for TFL1 expressing cells by isolating shoot apices from 42-day-old short-day-grown inflorescences just prior to onset of flower formation (**Fig. 1a**). Finally, we optimized low abundance ChIP by combining eight individual ChIP reactions per ChIP-seq replicate. We conducted FD ChIP-seq in analogous fashion using a published ^22^, biologically active (**Supplementary Fig. 1c**), genomic fusion protein (gFD-GUS *fd-1*). Our ChIP-seq uncovered 3,308 and 4,422 significant TFL1 and FD peaks (MACS2 summit qval≤10^-10^), respectively (**Fig. 1b**). The TFL1 peaks significantly overlapped with the FD peaks (72% overlap, pval<10^-300^, hypergeometric test; **Fig. 1b, c**). We performed *de novo* motif analysis and identified the bZIP G-box *cis* motif, a known FD binding site ^23^, as most significantly enriched (pval<10^-470^) and frequently present (> 84%) under TFL1 and TFL1/FD co-bound peaks (**Fig. 1d** and **Supplementary Figs. 2, 3**). To test whether TFL1 occupancy depends on presence of functional FD, we next performed TFL1 ChIP-seq in the *fd-1* null mutant.

**Figure 1.**
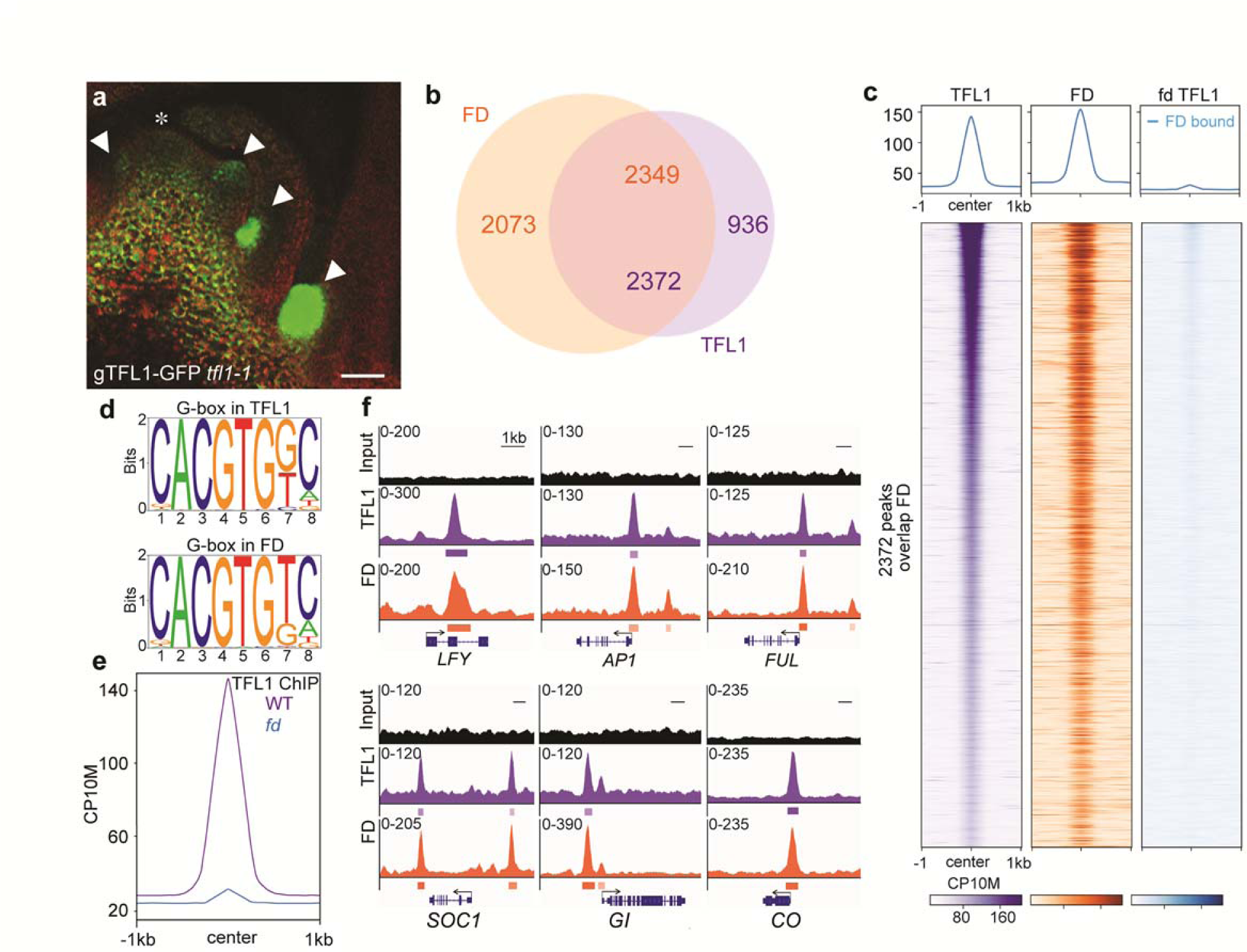
**TFL1 is recruited by FD to target loci. a**, gTFL1-GFP expression in branch meristems in the axils of cauline leaves (arrowheads) in 42-day-old short-day grown plants. Asterisk: shoot apex. **b**, Overlap of significant (MACS2 q value ≤10^-10^) FD and TFL1 ChIP-seq peaks. **c**, Heatmaps of TFL1, FD and TFL1(fd) ChIP-seq peaks centered on TFL1 binding peak summits, ranked from lowest to highest TFL1 summit q-value. **d**, Most significant motifs identified by *de novo* motif analysis under TFL1 or FD peak summits. **e**, TFL1 ChIP in the wild type (WT) compared to that in the *fd-1* null mutants. CP10M, counts per 10 million reads. **f**, Browser view of TFL1 and FD binding peaks at genes that promote onset of reproductive development ^1^ (*SOC1, GI, CO*) or switch to flower fate ^25^ (*LFY, AP1, FUL*). Significant peaks (summit qval≤10^-10^) according to MACS2 are marked by horizontal bars, with the color saturation proportional to the negative log 10 q value (as for the narrowPeak file format in ENCODE). See also **Supplementary Figs. 1-3** and **Supplementary Dataset 1**.

TFL1 chromatin occupancy was strongly reduced in *fd-1* (**Fig. 1c, e**). Our data suggest a prominent role of TFL1 in the nucleus and indicate that FD recruits TFL1 to the chromatin of target loci.

Annotating FD and TFL1 peaks to genes identified 2,699 joint TFL1 and FD targets. Gene Ontology (GO) term enrichment analysis implicates these targets in abiotic and endogenous stimulus response and reproductive development (**Supplementary Table 1**). Joint TFL1 and FD peaks were present at loci that promote onset of the reproductive phase in response to inductive photoperiod ^1, 3, 24^ like *GIGANTEA* (*GI*), *CONSTANS* (*CO*), and *SUPPRESSOR OF CONSTANS1* (*SOC1*) and of flower fate ^24, 25^ such as *LFY*, *APETALA1* (*AP1*), and *FRUITFULL* (*FUL*) (**Fig. 1f**). Identification of these TFL1 and FD bound targets fits well with TFL1’s known biological role (suppression of onset of reproduction and flower formation) and proposed molecular function (opposition of gene activation) ^4–6, 15^. We selected the *LFY* gene, which encodes a master regulator of flower fate (**Supplementary Fig. 4a-f**) ^6, 26–28^, to further probe the molecular mechanism of action of TFL1. To test whether *LFY* expression is directly repressed by the TFL1-FD complex, we generated transgenic plants expressing a steroid inducible version of TFL1 (TFL1^ER^; **Supplementary Fig. 5**). A single steroid treatment reduced *LFY* levels by 50% after 4 hours (**Supplementary Fig. 4g**), suggesting that the TFL1-FD complex represses *LFY*.

To better understand TFL1 recruitment to the *LFY* locus, we identified the genomic region sufficient and the cis motifs necessary for TFL1 association with *LFY*. TFL1 and FD peak summits were located in the second exon of *LFY* (**Fig.1f** and **Supplementary Fig. 4h, i)** and *LFY* reporters that lack the second exon were not repressed in response to TFL1 overexpression (**Supplementary Fig. 6a, b**). Exonic transcription factor binding sites, although rare, are found in both animals and plants, and frequently link to developmental regulation ^29, 30^. We identified three putative bZIP binding sites in *LFY* exon two: an evolutionarily conserved G-box and two partially conserved C-boxes (**Fig. 2a**). LFY exon two alone – when randomly integrated into the genome- was sufficient to recruit TFL1 (**Fig. 2b**). The recruitment was abolished when the three bZIP binding motifs were mutated (**Fig. 2b**). We confirmed recruitment of TFL1 to *LFY* exon two via the three bZIP binding motifs in a heterologous system (**Supplementary Fig. 6c**).

**Figure 2.**
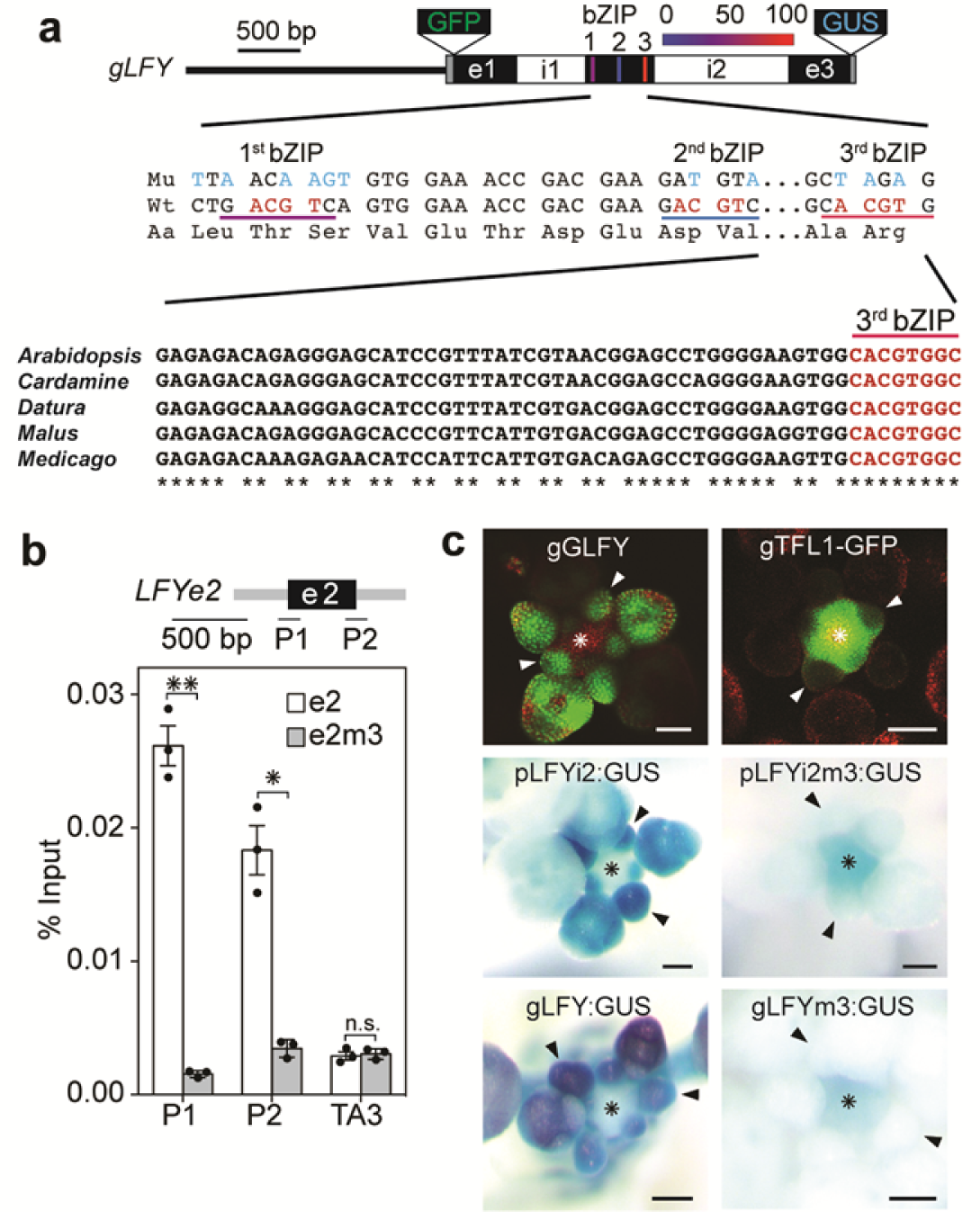
Exonic bZIP cis motifs mediate TFL1 recruitment to *LFY* **a**, Top: Fusion of genomic *LFY* construct with GFP (gGLFY) or beta-glucuronidase (gLFY-GUS). Location of putative bZIP binding motifs in *LFY* exon 2, color-coded based on conservation. For the pLFYi2:GUS reporter diagram see **Supplementary Fig. 5a**. Center: Mutation of the three bZIP binding motifs without changing the primary amino acid sequence. Bottom: Evolutionary conservation of the G-box. **b**, TFL1 recruitment to *LFY* exon 2 (e2) or a bZIP binding site mutated version thereof (e2m3) in 42-day old short-day-grown plants. Shown are mean ± SEM of three independent experiments (black dots). * *p*-value < 0.05, ** *p*-value < 0.01; unpaired one-tailed *t*-test. n.s.: not significant (*p*-value > 0.05). **c**, Top view of inflorescence apices with flower primordia. Plants were grown in long day. Top: Expression domain of LFY and TFL1 proteins. Center and Bottpm: Accumulation of beta-glucuronidase in a wild-type or bZIP binding site mutated (m3) reporter (pLFYi2:GUS) or genomic construct (gLFY-GUS). Arrowheads: flower primordia; asterisks: shoot apices; Scale bars: 2 mm. See also

We next assessed the contribution of the bZIP motifs to spatiotemporal *LFY* accumulation. Mutating the three bZIP binding sites in a *LFY* reporter that contains exon two (pLFYi2-GUS) or in a genomic *LFY* construct (gLFY-GUS) caused ectopic reporter expression in the center of the inflorescence shoot apex, the region where TFL1 protein accumulates during reproductive development (**Fig. 2c**) ^14^. Likewise, *LFY* is ectopically expressed in the inflorescence shoot apex of *tfl1* mutants during reproductive development ^6^. The combined data suggest that the TFL1-complex represses *LFY* via exonic bZIP motifs.

Surprisingly, the bZIP binding site mutations in the second exon of *LFY* in addition strongly reduced reporter expression in flower primordia (**Fig. 2c**), suggesting a role in *LFY* upregulation, possibly via FT. Based on prior studies ^24, 31–34^, *LFY* was not thought to be an immediate early FT target. While *ft* mutants display reduced *LFY* expression, this could be an indirect effect of delayed cessation of vegetative development ^35^. To assess whether FT promotes *LFY* expression, we therefore depleted FT specifically during the reproductive phase (**Fig. 3a**, **Supplementary Fig. 7a, b**). Onset of reproduction was normal in the conditional *ft* mutant (p4kbFT:amiRFT). However, p4kbFT:amiRFT significantly delayed flower formation and failed to upregulate *LFY* expression (**Fig. 3a** and **Supplementary Fig. 7**). In an orthogonal approach we induced endogenous *FT* expression by treating 42-day-old short-day grown plants with a single far-red enriched long-day photoperiod (FRP). FRP enhances *FT* induction by photoperiod ^36, 37^. FRP triggered significant *LFY* induction in the wild type, but not in the *ft* null mutant (**Supplementary Fig. 8**). *LFY* upregulation by FRP was furthermore dependent on presence of functional bZIP binding sites (**Fig. 3b**). We next generate a steroid inducible version of FT (FT-HA^ER^; **Supplementary Fig. 9a-c**). A single steroid treatment triggered significant *LFY* upregulation after 4 hrs (**Supplementary Fig. 9d, e**). LFY induction by FT-HA^ER^ was dependent on the presence of the bZIP motifs in the second exon (**Supplementary Fig. 9e**). The combined data reveal that FT activates *LFY* expression via the bZIP binding sites in the second exon.

**Figure 3.**
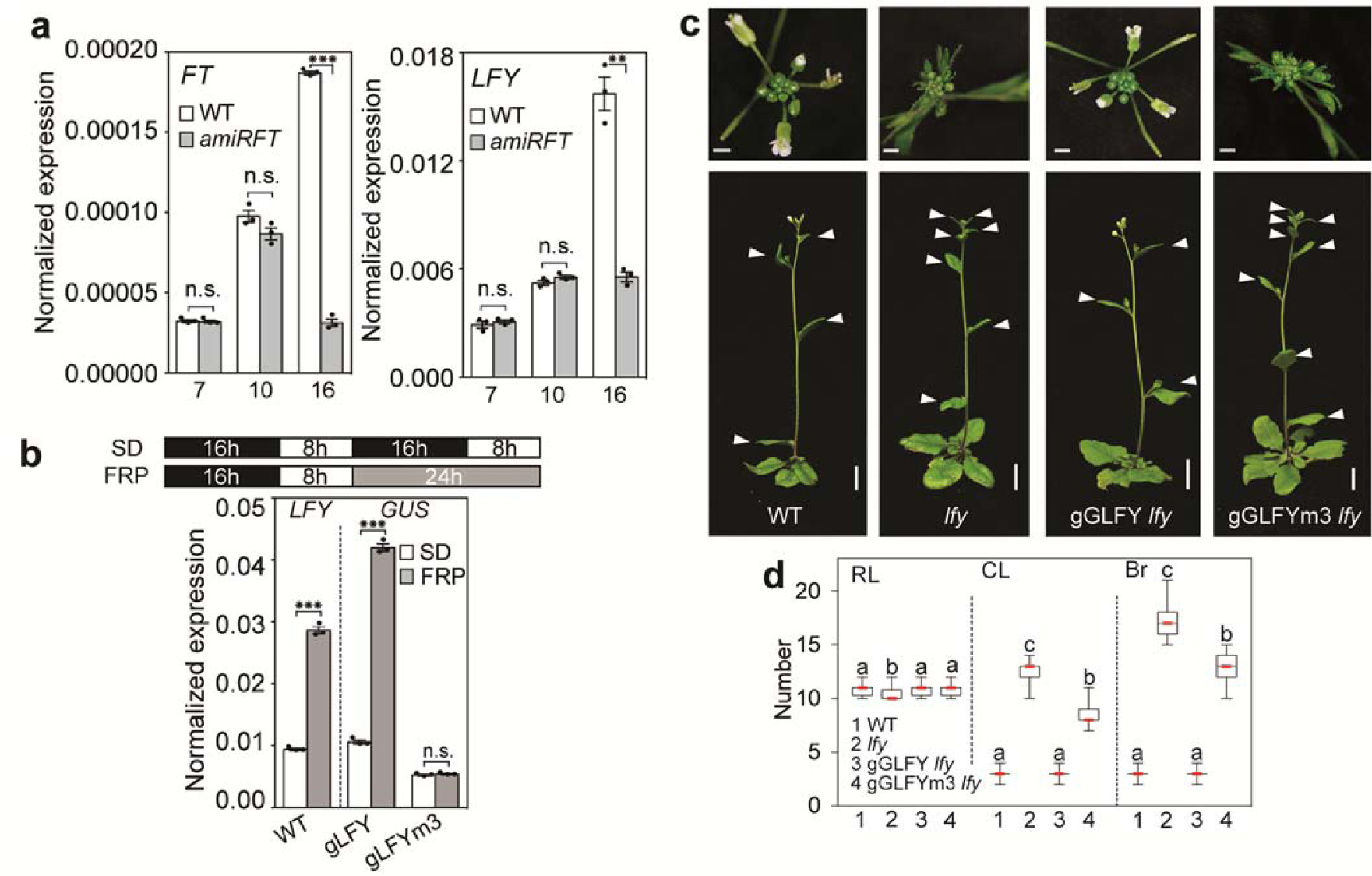
FT upregulates *LFY* expression to promote floral fate. a, Expression of *FT* and *LFY* prior to onset of reproductive development (day 7 and day 10) and during reproductive development (day 16) in long-day-grown wild type (WT) and in plants where *FT* was specifically downregulated during reproductive development (*pFT4kb:amiRFT*). **b**, Effect of a single far-red enriched photoperiod (FRP) on *LFY* (left) and reporter (GUS, right) accumulation in 42-day-old short day (SD) grown plants. (a, b) Expression was normalized over *UBQ10.* Shown are mean ± SEM of three independent experiments (black dots). unpaired one-tailed *t*-test; *** *p*-value < 0.001, ** *p*-value < 0.01; n.s.: not significant (*p*-value > 0.05). **c,** Rescue of *lfy-1* null mutants by genomic GFP-tagged LFY or a bZIP binding site mutated version thereof (gGLFYm3) in long-day-grown plants. Representative inflorescence images (top and side view). For a diagram of the mutations see Fig. 2a. **d**, Phenotype quantification. RL, rosette leaves; CL, cauline leaves; Br, branches. Box plot-median (red line; n = 15 plants), upper and lower quartiles (box edges), and minima and maxima (whiskers). Letters: significantly different groups *p*-value < 0.05 based on Kruskal-Wallis test with Dunn’s *post hoc* test. Scale bars, 1 cm. See also **Supplementary Figs. 6 - 10**.

We next probed the biological importance of the *LFY* bZIP binding sites for inflorescence architecture. While a genomic GFP-tagged *LFY* construct (gGLFY) fully rescued the *lfy-1* null mutant (14 out of 15 transgenic lines), a construct which preserves LFY protein sequence but has mutated bZIP binding sites yielded only partial rescue (gGLFYm3; 15 out of 15 transgenic lines) (**Fig. 3c, d**): onset of flower formation was significantly delayed in the gGLFYm3 *lfy-1* plants and *LFY* accumulation was very strongly reduced (**Fig. 3c, d** and **Supplementary Fig. 10**). The dramatic reduction of *LFY* accumulation is striking given the many additional inputs into *LFY* upregulation previously identified ^31, 33, 38, 39^. Our combined data uncover a pivotal role of FT in *LFY* upregulation and reveal that FT promotes flower formation via *LFY*. The bZIP mutations did not cause the terminal flower phenotype typical of *tfl1* mutants. This is expected since the bZIP mutations at the *LFY* locus mimic combined loss of TFL1 and FT activity, which likewise lacks a terminal flower phenotype ^40^.

Having identified *LFY* as a target under dual opposite regulation by TFL1 and FT, we next investigated the mechanism underlying their antagonism at this locus. To test for possible competition between FT and TFL1 at the chromatin, we conducted anti-HA ChIP qPCR in 42-day-old short-day grown FT-HA^ER^ gTFL1-GFP plants four hours after mock or steroid application. Estradiol induction led to rapid recruitment of FT-HA to the second exon of *LFY*, the region occupied by FD and TFL1 (compare **Fig. 4a** to **Supplementary Fig. 4h, i**). Anti-GFP ChIP qPCR performed on the same sample uncovered a concomitant reduction of TFL1 occupancy (**Fig. 4a**). Thus, FT upregulation competes TFL1 from the chromatin. To test whether endogenous FT can compete TFL1 from the *LFY* locus, we induced FT by a single FRP. Photoinduction of FT likewise significantly reduced TFL1 occupancy at the *LFY* chromatin (**Fig. 4b**). By contrast, photoinduction of FT did not alter FD occupancy at the *LFY* locus (**Fig. 4c**). That loss of TFL1 from the LFY locus is due to competition by FT is further supported by the finding that neither steroid nor FRP induction of FT reduced *TFL1* mRNA accumulation (**Supplementary Figs. 8c, 9d**). As observed for TFL1 (**Fig. 2b**), LFY exon two was sufficient and the bZIP sites in it necessary for FT-HA^ER^ recruitment (**Fig. 4d**). Finally, we examined whether FT induction by FRP competes TFL1 from the other direct TFL1-FD target loci identified (**Fig. 1f**). FRP treatment reduced TFL1 occupancy at the peak summits of all loci tested (**Fig. 4e**). Thus, one mechanism for the antagonism between FT and TFL1 ^15–17^ is through competition for FD bound at the chromatin of shared target loci to direct gene activation or repression, respectively (**Fig. 4f**).

**Figure 4.**
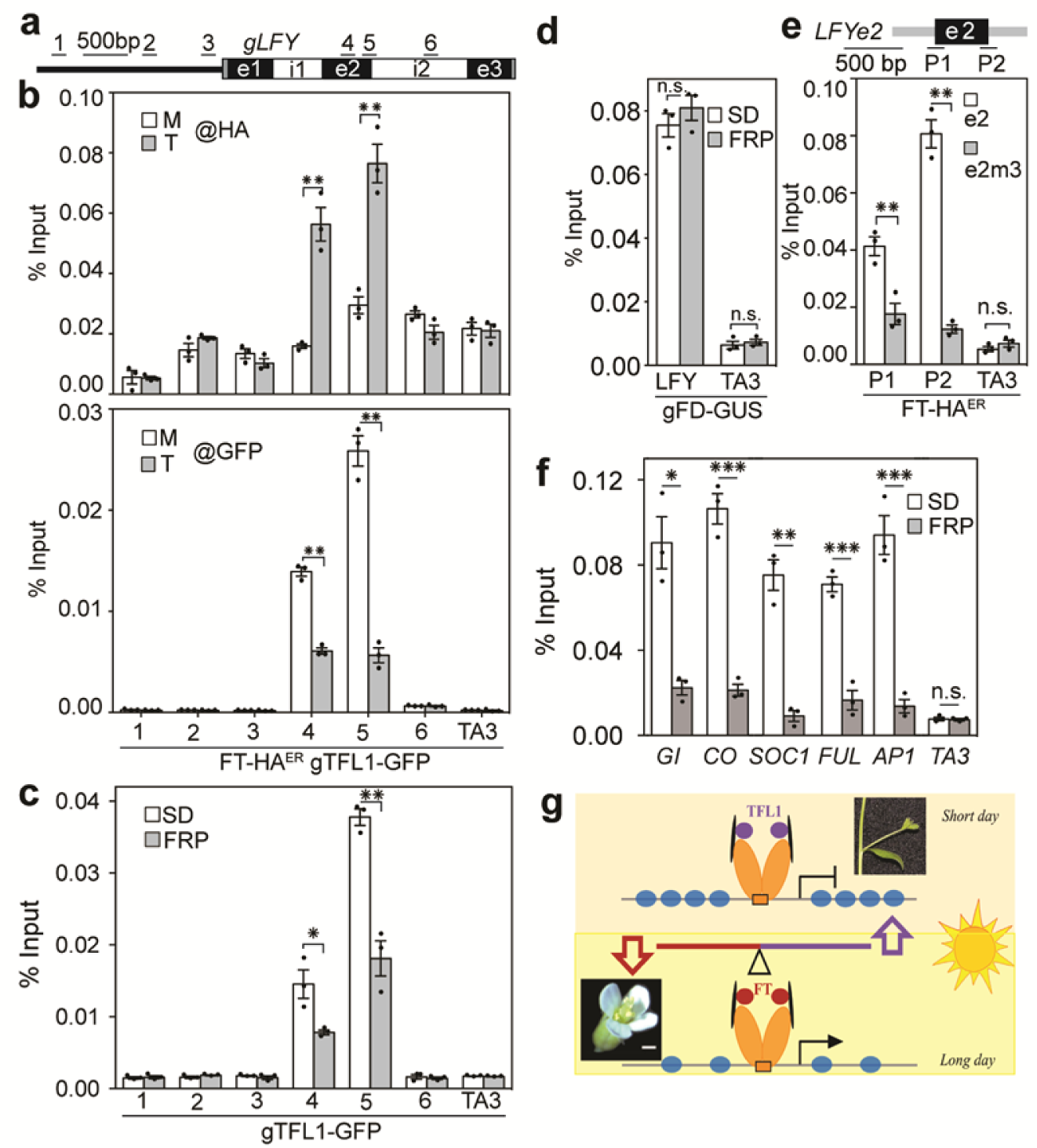
**FT competes TFL1 from the chromatin. a**, *LFY* locus and primers used. **b**, Top: FT-HA^ER^ occupancy at the *LFY* locus after four-hour mock (M) or steroid (T) treatment. Bottom: gTFL1 occupancy in the same sample. **c**, Effect of *FT* upregulation by photoperiod (FRP, 24 hours) on TFL1 occupancy at the *LFY* locus. **d**, Effect of *FT* upregulation by photoperiod (FRP, 24hours) on FD occupancy at the *LFY* locus. **e**, FT-HA^ER^ recruitment after four-hour steroid treatment to *LFY* exon 2 (e2) or a bZIP binding site mutated version thereof (e2m3). **f**, Effect of FT upregulation by photoperiod (FRP) on TFL1 occupancy at TFL1-FD target loci identified in Fig. 1f. (b - f) ChIP was performed in 42-day-old short-day-grown plants. Shown are mean ± SEM of three independent experiments (black dots). * *p*-value < 0.05, ** *p*-value < 0.01; *** *p*-value < 0.001; unpaired one-tailed *t*-test. n.s.: not significant (*p*-value > 0.05). **g,** Model for antagonist roles of TFL1 (purple circles) and FT (red circles) in promoting branch fate or floral fate, respectively. Increased FT accumulation leads to competition of TFL1 from bZIP transcription factor FD bound to chromatin and to onset of flower formation. FD dimers (orange ovals), 14-3-3 proteins (black disks).

Our data places florigens directly upstream of LFY, yet prior genetic data suggest that florigens act both upstream of and in parallel with LFY ^32, 41^. To gain insight into the gene expression programs repressed by the TFL1-FD complex, we next conducted RNA-seq with and without FRP. We isolated inflorescences with associated primordia from 42-old short day-grown *ft* mutant, wild-type and *tfl1* mutant plants. After FRP treatment, we identified the significant gene expression changes in each genotype relative to untreated siblings (**Supplementary Fig. 11)**. In particular, we were interested in TFL1-FD complex bound loci that exhibit FT-dependent de-repression upon photoinduction. 604 TFL1-FD bound genes were significantly (DESeq2 adjusted p<0.005) de-repressed upon FRP treatment in the wild type or in *tfl1* mutants but not in *ft* mutants (**Fig. 5a**).

**Figure 5.**
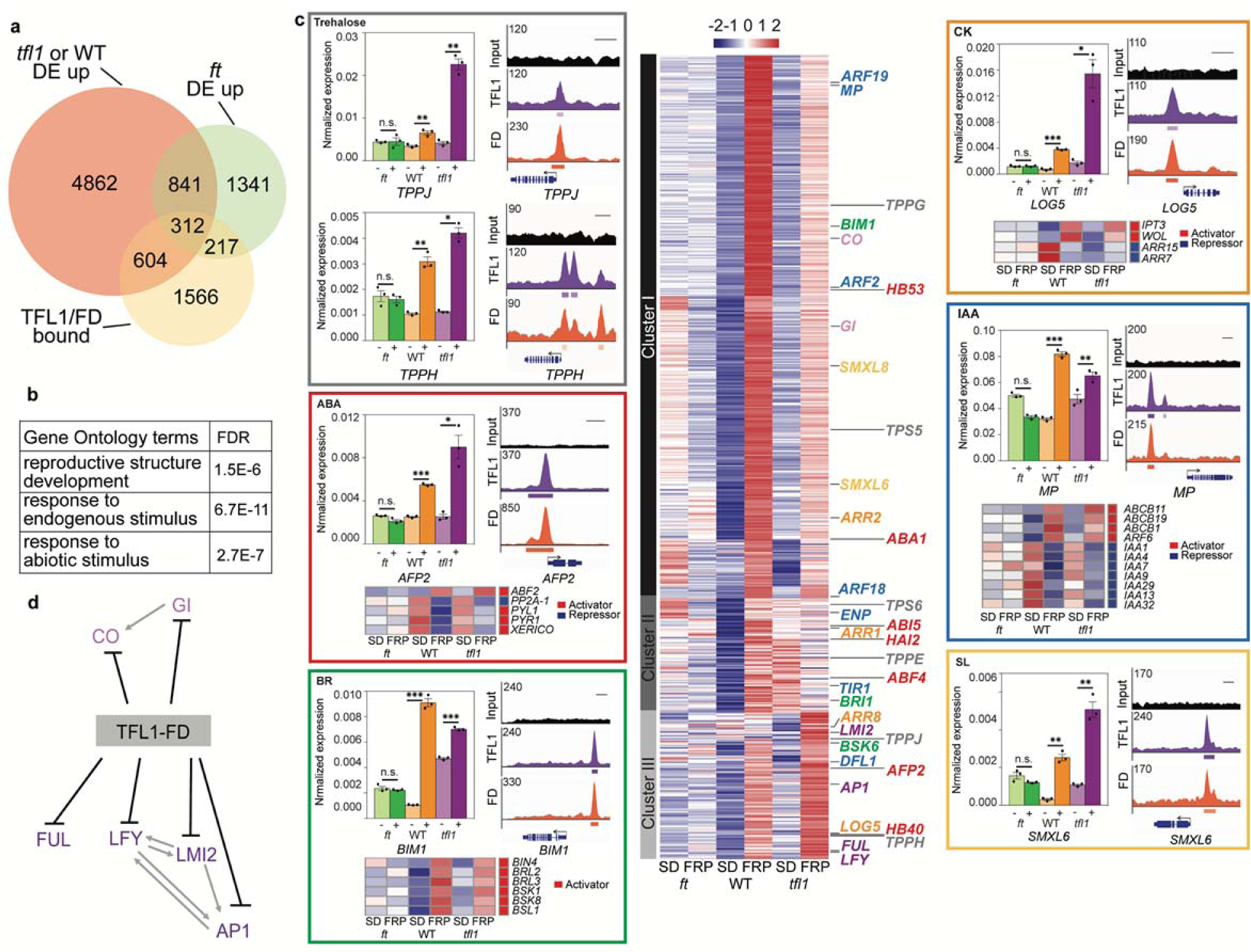
**Genes directly repressed by the TFL1-FD complex. a**, 604 genes bound by TFL1 and FD are de-repressed after a single FRP specifically in the wild type or in *tfl1* mutants. **b**, Gene ontology term enrichment (Goslim AgriGO v2) for the 604 direct TFL1-FD complex repressed genes. **c**, K-means clustering of the direct TFL1-FD complex repressed genes (center). Heatmap and gene names colour coded for the pathway they act in. Onset of reproduction (light purple), floral fate (purple), sugar signalling (grey), abscisic acid (ABA, red), brassinosteroid (BR, green), cytokinin (CK, orange), auxin (IAA, blue) and strigolactone (SL, yellow). Color coded boxes: independent confirmation of gene expression changes of direct TFL1-FD targets and ChIP-seq binding screenshots. Heatmaps below indirect (downstream) FT-dependent gene expression changes of known pathway activators (red in sidebar) or repressors (blue in sidebar) indicative of de-repression of the hormonal pathway. Auxin: upregulation a positive auxin response regulator (*ARF6*) and downregulation of negative auxin response regulators (*IAAs*) ^55^. Brassinosteroid: upregulation of multiple positive signalling pathway components (*BRL2*, *BRL3*, BSK1, *BSK8*, *BSL1* and *BIN4*) ^56^. Cytokinin: upregulation of cytokinin production (*IPT3*) and perception (*WOL*) and downregulation of negative cytokinin response regulators (*ARR7, ARR15*) ^57^. Abscisic acid: Upregulation of some positive regulators of abscisic acid response (*ABF2*), while that of others was decreased (*PYL1*, *PYR1*, *XERICO*) ^58^. * *p*-value < 0.05; ** *p*-value < 0.01, *** *p*-value < 0.001; unpaired one-tailed *t*-test in. n.s.: not significant (*p*-value > 0.05). **d**, Interaction network of TFL1-FD complex repressed targets linked to onset of reproduction (CO, GI) and floral fate (all others). Previously described interactions ^1, 3, 25^ are indicated by grey arrows. See also **Supplementary Figs. 11 - 14**.

Gene ontology term enrichment linked the TFL1-FD repressed genes to reproductive development and to response to endogenous and abiotic signals (**Fig. 5b**). We performed k-means clustering of the 604 genes and identified three main patterns of gene expression. Genes encoding promoters of floral fate ^25^ (*LFY*, *AP1*, *FUL* and *LMI2*) clustered together (cluster III in **Fig. 5c**) and displayed stronger upregulation in *tfl1* mutants than in the wild type. *SOC1* clusters with these genes, but was not included in further analyses because it was weakly, but significantly, de-repressed in *ft* mutants (**Supplementary dataset 1** and **Supplementary Fig. 12**). By contrast, *CO* and *GI,* which promote cessation of vegetative development ^1, 3^, were more strongly upregulated in the wild type than in *tfl1* mutants (cluster I in **Fig. 5c**), perhaps because these genes are already partially de-repressed in *tfl1* mutants in the absence of FRP treatment.

Indeed, both RNA-seq and qRT-PCR of independent biological replicates revealed higher accumulation of GI and CO in untreated tfl1 mutant compared to wild-type plants (**Supplementary dataset 1** and **Supplementary Fig. 12**). Cluster II genes are only upregulated in the wildtype and may represent genes de-repressed transiently. We confirmed FT-dependent de-repression of the regulators of reproductive development and floral fate using independent biological samples (**Supplementary Fig. 12a, b**). Thus, the TFL-FD complex directly represses key regulators that trigger onset of reproductive development and flower fate (**Fig. 5c**).

In addition, our combined ChIP-seq and RNA-seq analysis identified novel direct TFL1-FD repressed targets. These include components of sugar signalling (trehalose 6-phosphate phosphatases (TPPs)) and genes linked to hormone response (abscisic acid, cytokinin, brassinosteroid, auxin and strigolactone) (**Fig. 5c, d**). TFL1 and FD strongly bound select genes in these signalling pathways and qRT-PCR of independent biological samples confirmed their FT-dependent upregulation by FRP photoinduction (**Fig. 5c**). To probe for a role of TFL1-FD in repression of hormone signalling and response, we assessed indirect, downstream, gene expression changes by identifying FT-dependent genes expression changes. We identified genes not bound by TFL1 or FD yet significantly de-repressed (DESeq2 adjusted p<0.005) in the wild type and in *tfl1*, but not in *ft*. This revealed de-repression of the auxin, brassinosteroid and cytokinin hormone pathways (**Fig. 5c**). The effect on abscisic acid signalling was more complex (**Fig. 5c**). Altogether, the downstream gene expression changes support a role for the TFL1-FD complex in repression of diverse hormone pathways. Finally, we identified transcriptional co-regulators and chromatin regulators among the direct TFL1-FD repressed targets (**Supplementary Fig. 13a, b**). These proteins may assist in cell fate reprogramming during onset of flower formation.

Some of the identified signalling pathway components may contribute to the switch to floral fate. For example, auxin and the auxin-activated transcription factor MONOPTEROS (MP) promote floral fate in *Arabidopsis* ^42^, while the tomato ortholog of TFL1 executes its role in inflorescence architecture at least in part by modulating auxin flux and response ^43^. While the TPPs and brassinosteroid signalling have not yet been implicated in promoting floral fate in *Arabidopsis*, they oppose branching in Maize ^44^ and Setaria ^45^ inflorescences, respectively. In addition, components of strigolactone, cytokinin, auxin, abscisic acid as well as sugar signalling may promote branch outgrowth ^46, 47^. Further support for a possible role of these signalling pathway components in branch outgrowth comes from phenotypic analyses. A single FRP reprogrammed branches to flowers subtended by cauline leaves in the wild type (see also Ref ^48^) and more strongly, in *tfl1* (**Supplementary Fig. 14a - j**). It also triggered a significant increase in primary branch outgrowth in both wild type and *tfl1* (**Supplementary Fig. 14k**). FRP had no phenotypic effect in *ft* mutants. Our combined data suggest that florigens tune plant form to the environment by controlling expression of master developmental regulators, endogenous signalling pathway components and chromatin remodelling and modifying proteins.

Here identify *LFY*, a master regulator of flower fate ^27, 28^, as a target under dual opposite transcriptional regulation by TFL1 and FT and show that FT activates *LFY* to promote floral fate in axillary meristems during inductive photoperiod. We provide a molecular framework for the antagonistic roles of the florigen family members FT and TFL, which relies on competition for bZIP transcription factor mediated access to binding sites at regulatory regions of shared target loci. We find that TFL1 does not merely prevent access of the FT co-activator to the chromatin but is an active repressor, as mutating florigen recruitment sites resulted in *LFY* de-repression in the TFL1 expression domain. We identify key regulators of onset of the reproductive phase and of floral fate as immediate early TFL1-FD complex repressed gene. In addition, we link the TFL1-FD complex to direct repression of multiple hormone signalling pathways and chromatin regulators. In combination, our data reveal first insight into how TFL1 and FT alter the developmental trajectory of primordia in response to seasonal cues and sculpt inflorescence architecture for enhanced reproductive success. Our analyses set the stage for elucidating communalities as well as differences between florigen regulated cell fate reprogramming during flower initiation and other traits under seasonal control by florigens such as tuberization, bulb formation and seed dormancy ^49–52^. Perturbation of the relative balance of opposing florigen activities underlies domestication of wild species, modulating desirable traits like decreased juvenility, everbearing and determinate growth habit ^12, 49, 53, 54^. Hence mechanistic insight into the activities and targets of florigens will also benefit traditional or genome editing based crop improvement.

## URLs of R packages

DESeq2: https://bioconductor.org/packages/release/bioc/html/DESeq2.html ggplot2: https://cran.r-project.org/web/packages/ggplot2/index.html pheatmap: https://cran.r-project.org/web/packages/pheatmap/index.html RColorBrewer: https://cran.r-project.org/web/packages/RColorBrewer/index.html PCC analysis: https://stat.ethz.ch/R-manual/R-devel/library/stats/html/cor.html Distance calculation: https://www.rdocumentation.org/packages/stats/versions/3.6.1/topics/dist kmeans: https://www.rdocumentation.org/packages/stats/versions/3.6.1/topics/kmeans.

## Methods

Methods, including statements of data availability and any associated accession codes and references, are available in the online version of the paper. Note: Any Supplementary Information and Source Data files are available in the online version of the paper.

## Acknowledgements

We thank undergraduate students Gabriela M. Blandino, Xindi Chen and Zubaida Salman and Dietrich James Nigh for help with the experiments, Dr. John D. Wagner for input on the manuscript and the University of Pennsylvania Plant Biology group as well as Wagner lab members for feedback on this project. This research was funded by National Science Foundation IOS grant 1557529 and 1905062 to DW. The ChIP-seq and RNA-seq data was deposited to the GEO database (GSE141894).

## Author contributions

D.W. and Y.Z. conceived of the study and Y.Z. conducted the majority of the experiments. S.K. conducted the bioinformatic analyses, N.Y. and C.W.J. identified the optimal stage to study axillary meristem regulation by TFL1, conducted initial TFL1 ChIP analyses and mapped the FD binding sites in *LFY*. R.J. and Y.Z. constructed and sequenced ChIP-seq libraries, K.G generated the biologically active genomic GFP-TFL1 construct. D.W. wrote the paper with input from all authors.

## Materials and Methods

### Plant materials

*Arabidopsis* ecotype Columbia plants were grown in soil at 22°C in long-day photoperiod (LD, 16h light / 8h dark, 100 µmol/m2 s) or short-day photoperiod (SD, 8h light / 16h dark, 120 µmol/m2 s). gFD-GUS^1^, *fd-1* null mutants^2^, *tfl1-14* hypomorph mutant^3, 4^, *tfl1-1* severe mutant^3–5^, *lfy-1* null mutant^6–8^, *ft-10* null mutants ^9^, 35S:LFY^10^ and 35S:TFL1^11^ were previously described. 35S:LFY (Landsberg *erecta)* was introgressed into the Columbia background through backcrossing. *gFD-GUS* was crossed into the *fd-1* null mutant background.

### Constructs for transgenic plants

For *gTFL1-GFP*, GFP followed by a peptide linker (GGGLQ) was fused to an 8.4 kb BamHI (NEB, R0136S) genomic fragment from lambda TFG4 (Ref.^12^). This fragment was introduced into the binary vector *pCGN1547* (Ref.^13^). For *TFL1^ER^* and *FT*-*HA^ER^*, *TFL1* and *FT* were PCR amplified from cDNA; in the case of FT, the 3’ primer contained three times Hemagglutinin (HA) plus a stop codon. PCR products were cloned into *pENTRD-TOPO* (Invitrogen, K243520) and shuffled into *pMDC7* (Ref.^14^) by LR reaction (Invitrogen, 11791-020).

For *LFY-GUS* reporters, the bacterial beta-glucuronidase (*GUS*) gene from *pGWB3* (Ref.15) binary vector was fused with *pENTRD*-*TOPO* vector containing the 2,290-bp *LFY* promoter ^16^ alone (pLFY:*GUS*), the *LFY* promoter and *LFY* genic region up to and including the first intron (pLFYi1:GUS), or the *LFY* promoter and *LFY* genic region up to and including the second *LFY* intron (*pLFYi2*:GUS) by LR reaction. bZIP binding site mutations in *LFY* exon 2 (*pLFYi2m3*:GUS) were generated by Ω-PCR ^17^.

*gGLFY* was constructed by PCR amplifying a 4,929-bp genomic *LFY* fragment (*gLFY*), including the 2,290-bp *LFY* promoter, from genomic DNA followed by cloning into a KpnI-HF (NEB, R3124S) and NotI-HF (NEB, R3189S) digested *pENTR3C* by Gibson Assembly. Next, GFP was inserted at position + 94 bp, as previously described for *pLFY:GLFY* ^18, 19^, by Ω-PCR^17^. bZIP binding sites mutations were introduced into *pENTR3C*-*gGLFY* by Ω-PCR to generate *gGLFYm3*. Both constructs were shuffled into *pMCS:GW* ^20^ using LR reaction. To create *gLFY*:*GUS*, the 4,929-bp genomic *LFY* clone minus stop codon and GUS fragments were PCR amplified and inserted into linearized *pENTR3C* by Gibson Assembly. For *gLFYm3*:*GUS*, bZIP binding site mutations were introduced into *pENTR3C-gLFY:GUS* by Ω-PCR. To test recruitment of TFL1 and FT to exon 2 of *LFY*, wild-type (e2) or bZIP binding site mutated exon 2 (e2m3) were PCR amplified and cloned into *pGWB3* (Ref. ^15^).

For *pFT4kb:amiRFT*, a 3,994-bp truncated *FT* promoter ^21^ was PCR amplified from genomic DNA as was the published *amiRFT* ^22^ from *pRS300* (Ref. ^23^). The amiRFT fragments were introduced into EcoRI-HF (NEB, R3101S) digested *pENTR3C* (Thermo Fisher Scientific, A10464) by Gibson Assembly (NEB, E5510S) and shuffled into binary vector *pMCS:GW* ^20^ using LR reaction, which resulted in pMCS:amiRFT. The previously described 3994-bp *FT* promoter was amplified and inserted to XhoI (NEB, R0146S) digested pMCS:amiRFT by Gibson Assembly.

Genomic DNA was extracted using the GenElute Plant Genomic DNA Miniprep Kit (Sigma-Aldrich, G2N70). Primer sequences are listed in Supplementary Table 2. All constructs were sequence verified prior to transformation into plants with Agrobacterium strain GV3101 by floral dip ^24^. Plant lines generated are listed in Supplementary Table 3.

### Imaging

Images were taken with a Canon EOS Rebel T5 camera for plant phenotypes and yeast one hybrid assays, or with a stereo microscope (Olympus SZX12) equipped with a color camera (Olympus LC30) for GUS images and inflorescence phenotypes.

For GFP images, a Leica TCS SP8 Multiphoton Confocal with a 20X objective was used with a 488 nm excitation laser and emission spectrum between 520 to 550 nm (GFP) or 650 to 700 nm (chlorophyll autofluorescence). Imaging was conducted essentially as previously described ^25, 26^, except for *tfl1-1* gTFL1-GFP in short day photoperiod, where shoot apices were sectioned longitudinally on an oscillating tissue slicer (Electron Microscopy Sciences, OTS-4000) after embedding in 5% Agar (Fisher Scientific, DF0812-07-1).

### Plant treatment and gene expression analysis

For test of gene expression, 16-day-old TFL1^ER^ or 12-day-old FT-HA^ER^ plants grown in LD were induced by a single spray application of 10 μmol beta-estradiol (Sigma-Aldrich, 8875-250MG) dissolved in DMSO (Fisher Scientific, BP231-1L) and 0.015% Silwet L-77 (PlantMedia, 30630216-3). Mock solution consisted of 0.1 % DMSO and 0.015 % Silwet. To probe FT recruitment to and TFL1 occupancy at the *LFY* locus, 42-day-old FT-HA^ER^ gTFL1-GFP plants grown in SD were treated by a single spray application of 10 μmol beta-estradiol or mock solution. In all cases, tissues were harvested 4 hours after treatment. To test for gain-of-function phenotypes in long-day photoperiod, FT-HA^ER^, TFL1^ER^ and FT-HA^ER^ gTFL1-GFP plants were treated with 10 μmol beta-estradiol or mock solution from 5-day onwards every other day until bolting.

Far-red light enriched photoinduction (FRP) was applied at the end of the short day (ZT8) using a Percival Scientific E30LED ^27^ with red (660nm) to far-red (730 nm) ratio = 0.5 and light intensity 80 µmol/m2 s. Control plants were kept in regular short-day conditions (16-hour dark and 8-hour light, red to far-red ratio = 12 and 120 µmol/m2s light intensity) for 24 hours. Light intensity and spectral composition were measured by an Analytical Spectral Devices FieldSpec Pro spectrophotometer.

For qRT-PCR analysis, total RNA was extracted from leaves or shoot apices using TRIzol (Thermo Fisher Scientific) and purified with the RNeasy Mini Kit (Qiagen, 74104). cDNA was synthesized using SuperScript III First-Strand Synthesis (Invitrogen, 18080051) from 1 μg of RNA. Real time PCR was conducted using a cDNA standard curve. Normalized expression levels were calculated using the 2^(-delta delta CT) method with the housekeeping gene *UBQ10* (AT4G05320) as the control. Where expression of multiple different genes was compared, normalized gene expression is shown relative to the control treatment. Primer sequences are listed in Supplementary Table 2.

### Yeast one-hybrid assay

LFY exon 2 (e2) and the bZIP binding site mutated version (e2m3) were cloned into the KpnI-HF and XhoI linearized pAbAi vector (Takara) by Gibson Assembly and integrated into the yeast genome following the Matchmaker Gold Yeast One-Hybrid protocol (Takara) and the Y1Golden strain (Takara). Coding sequences of FD and TFL1 were cloned into the pENTRD-TOPO vector. After sequencing, constructs were shuffled into either pDEST32 or pDEST22 (Takara) by LR reaction and transformed into the DNA binding region containing yeast strain. Empty pDEST32 and pDEST22 served as negative controls. Growth was assayed after serial dilution on growth media with or without 60 ng /ml Aureobasidin A (Clontech, 630499). Primer sequences are listed in Supplementary Table 2.

### ChIP-qPCR, ChIP-seq and data analysis

42-day-old, short-day grown plants were trimmed and 1.6 gram of non-bolted inflorescences were harvested from 36 plants. Chromatin immunoprecipitation was conducted as previously described ^28^ for ChIP-qPCR. For ChIP-seq, each biological replicate consisted of 8 individual IP reactions pooled into one MinElute PCR (Qiagen, 28004) purification column. For ChIP-seq and ChIP-qPCR, anti-GFP antibody (Thermo Fisher Scientific, A-11122; 1:200 dilution) and anti-GUS antibody (Abcam, ab50148; 1:200 dilution) were used. The antibodies were validated by the manufacturers. ChIP-qPCR was performed using Platinum Taq DNA Polymerase (Invitrogen, 10966034) and EvaGreen dye (Biotium, 31000). For ChIP-qPCR, the value of the ChIP samples was normalized over that of input DNA as previously described ^28^. Non-transgenic wild-type plants were used as the negative genetic control for anti-GFP and anti-GUS antibody ChIP. The *TA3* retrotransposon (AT1G37110) was used as the negative control region for ChIP-qPCR. Primer sequences are listed in Supplementary Table 2.

Anti-GFP ChIP-seq was performed for gTFL1-GFP (A), gTFL1-GFP (B), *fd-1* gTFL1-GFP and control samples (non-epitope containing plants). Anti-GUS ChIP-seq was performed for for *gFD-GUS*. Two biological replicates were sequenced in each case. Dual index libraries were prepared for the ten ChIP samples listed above and for four input samples using the SMARTer ThruPLEX DNA-Seq Kit (Takara Bio, R400406). Library quantification was performed with the NEBNext Library Quant Kit for Illumina (NEB, E7630). Single-end sequencing was conducted using High Output Kit v2.5 (Illumina, TG-160-2005) on the NextSeq500 platform (Illumina).

FastQC v0.11.5 was performed on both the raw and trimmed ^29^ reads (TRIMMOMATIC v0.36 ^29^ (ILLUMINACLIP:adapters.fasta:2:30:10 LEADING:3 TRAILING:3 MINLEN:50) to confirm sequencing quality ^30^. Reads with MAPQ ≥ 30 (SAMtools v1.7) ^31^ uniquely mapping to the Arabidopsis Information Resource version 10 (TAIR 10) ^32^ that were not flagged as PCR or optical duplicates by Bowtie2 v2.3.1(Ref. ^33 34^) were analysed further by principal component analysis using the plotPCA function of deepTools ^35^. Reads were further processed following ENCODE guidelines ^36^, followed by cross-correlation analysis with the predict function of MACS2 (Ref. ^37^) to empirically determine the fragment length. Significant ChIP peaks and summits (summit q-value ≤10^-10^) were identified in MACS2 relative to the pooled negative controls (ChIP in nontransgenic wild type); a total of 3,308 peaks for TFL1 and 4,422 for FD. Peak overlap (≥ 1 bp) was computed using BEDTools intersect v2.26.0 (Ref. ^38^) and statistical significance was computed using the hypergeometric test and assuming a ‘universe’ of 10,000 possible peaks ^39^. Heatmaps were generated using deepTools v3.1.2 (Ref. ^35^). The 3,308 TFL1 and 4,422 FD peaks were mapped to 3,699 and 4,493 Araport11 (Ref. ^40^) annotated genes, respectively, if the peak summit was intragenic or located ≤ 4 kb upstream of the transcription start site. Genomic distribution of peak summits were called using the ChIPpeakAnno library ^40, 41^.

*De novo* motif analysis was conducted using MEME-ChIP v5.0.2 (Discriminative Mode) ^42^ and HOMER v4.10 (Ref. ^43^) for MACS2 q-value ≥ 10^-10^ peaks (+/-250 base pairs) compared to genome matched background (unbound regions from similar genomic locations as the peak summits as previously described ^44, 45^.

Gene Ontology (GO) term enrichment analyses were performed using GOSlim in agriGO v2.0 (Ref. ^46^) and significantly enriched GO terms with q value <0.0001 (FDR, Benjamini and Yekutieli method ^47^) were identified.

### Correlation analyses

Public ChIP-seq datasets for FD ^48^ (European Nucleotide Archive accession PRJEB24874), LFY ^49^ and an unpublished LFY ChIP-seq dataset from our lab (GEO accession will receive shortly) were analysed as described above for TFL1 and FD ChIP-seq. To compare the relationship between all ChIP seq datasets we calculated Pearson correlation coefficients for reads in regions of interest using deeptools ^35^. Regions of interest were comprised of the combined significant peak regions (MACS2 ≤ q value 10^-10^) of all ChIP-seq datsets and read signal was derived from the sequencing-depth normalized bigwig file for each sample.

### RNA-seq and data analysis

A single 24-hour far-red enriched photoperiod (FRP) was applied to 42-day-old short-day grown *ft-10* mutants, wild type and *tfl1-1* mutants, starting at the end of the day (ZT8). After the treatment, 0.1 gram of inflorescence shoots were harvested for each biological replicate after removing all leaves and roots. Three biological replicates were prepared for each experiment. RNA quantity and quality were analysed by Qubit BR assay (Thermo Fisher Scientific, Q10210) and Agilent RNA 6000 Nano Kit (Agilent, 5067-1511) on an Agilent 2100 bioanalyzer, respectively. Libraries were constructed from1 ug total RNA using the TruSeq RNA Sample Prep Kit (Illumina, RS-122-2001). After library quantification with the NEBNext Library Quant Kit for Illumina (NEB, E7630), single-end sequencing was conducted using the NextSeq 500 platform (Illumina).

RNA-seq analysis was conducted using FastQC v0.11.5 (Ref. ^30^) on raw sequences before and after trimming using Trimmomatic v0.36 (Ref. ^29^) (ILLUMINACLIP:adapters.fasta:2:30:10 LEADING:3 TRAILING:3 MINLEN:50) to confirm sequencing quality ^30^.Reads were mapped using the STAR mapping algorithm ^50^ (--sjdbOverhang 100 --outSAMprimaryFlag AllBestScore --outSJfilterCountTotalMin 10 5 5 5 --outSAMstrandField intronMotif --outFilterIntronMotifs RemoveNoncanonical -- alignIntronMin 60 --alignIntronMax 6000 --outFilterMismatchNmax 2), to the TAIR10 Arabidopsis genome-assembly ^32^, and Araport11 Arabidopsis genome-annotation ^40^. Specific read coverage was assessed with HT-Seq ^51^ (--stranded=’no’ --minaqual=30).

For Principal Components Analysis (PCA), raw read counts were subjected to variance stabilizing transformation (VST) and projected into two principal components with the highest variance ^52–54^. In parallel, raw reads were adjusted for library size by DESeq2 (Ref. ^55^). Gene normalized z-scores were used k-means (MacQueen) clustering ^56^. After PCA, two biological replicates per genotype and treatment were selected for further analysis. Pairwise differential expression analyses were performed by comparing FRP and untreated pooled normalized read counts in each genotype using default DESeq2 parameters with no shrinkage and an adjusted p-value cut-off ≤ 0.005 (Ref. ^55^).

### Photoperiod shift phenotype analysis

A single 24-hour far-red enriched photoperiod shift (FRP) was applied to 42-day-old short-day grown *ft-10* mutants, wild type and *tfl1-1* mutants as for RNA-seq, followed by further growth in short day conditions. To asses onset of reproductive development, the number of rosette leaves formed were counted at bolting. To analyse the inflorescence architecture, the number of sessile buds, outgrowing branches, flower branches, and single flowers subtended by a cauline leaf were counted weekly after bolting until the first normal flower (not subtended by a cauline leaf) formed.

### Statistical analyses

The Kolmogorov-Smirnov (K-S) test ^57^ was used to assess whether the data were normally distributed. All ChIP and qRT-PCR data were normally distributed. An unpaired one-tailed *t*-test was used to test for changes in one direction. Error bars represent the standard error of the mean (SEM). Two to three independent biological replicates were analyzed. For multiple-group comparisons (phenotypes) the non-parametric Kruskal-Wallis test ^58^ followed by the Dunn’s *post hoc* test ^59^ were employed. Box and whisker plots display minima and maxima (whiskers), lower and upper quartile (box) and median (red vertical line).

## Supplementary figures

**Supplementary Figure 1.**
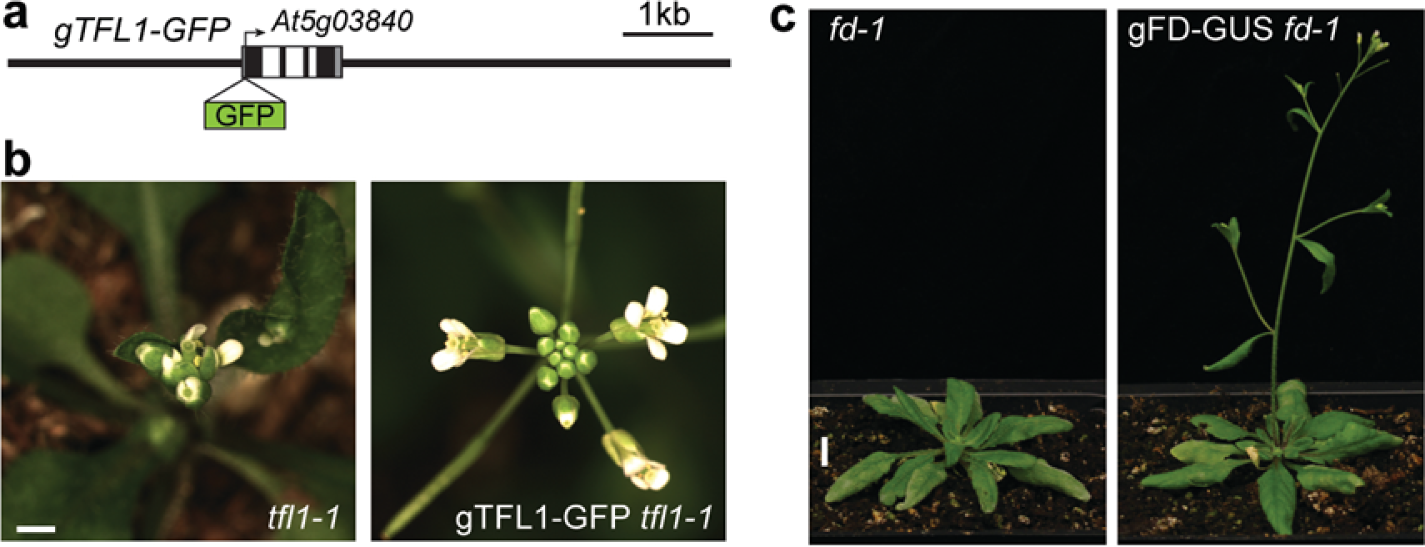
Biological activity of gTFL1-GFP and gFD-GUS. a, gTFL1-GFP construct. **b**, Rescue of the terminal flower phenotype of the severe *tfl1-1* mutant by gTFL1-GFP. Scale bar, 5 mm. **c**, Rescue of the late-flowering phenotype of the null *fd-1* mutant by gFD-GUS^18^. Scale bar, 1cm.

**Supplementary Figure 2.**
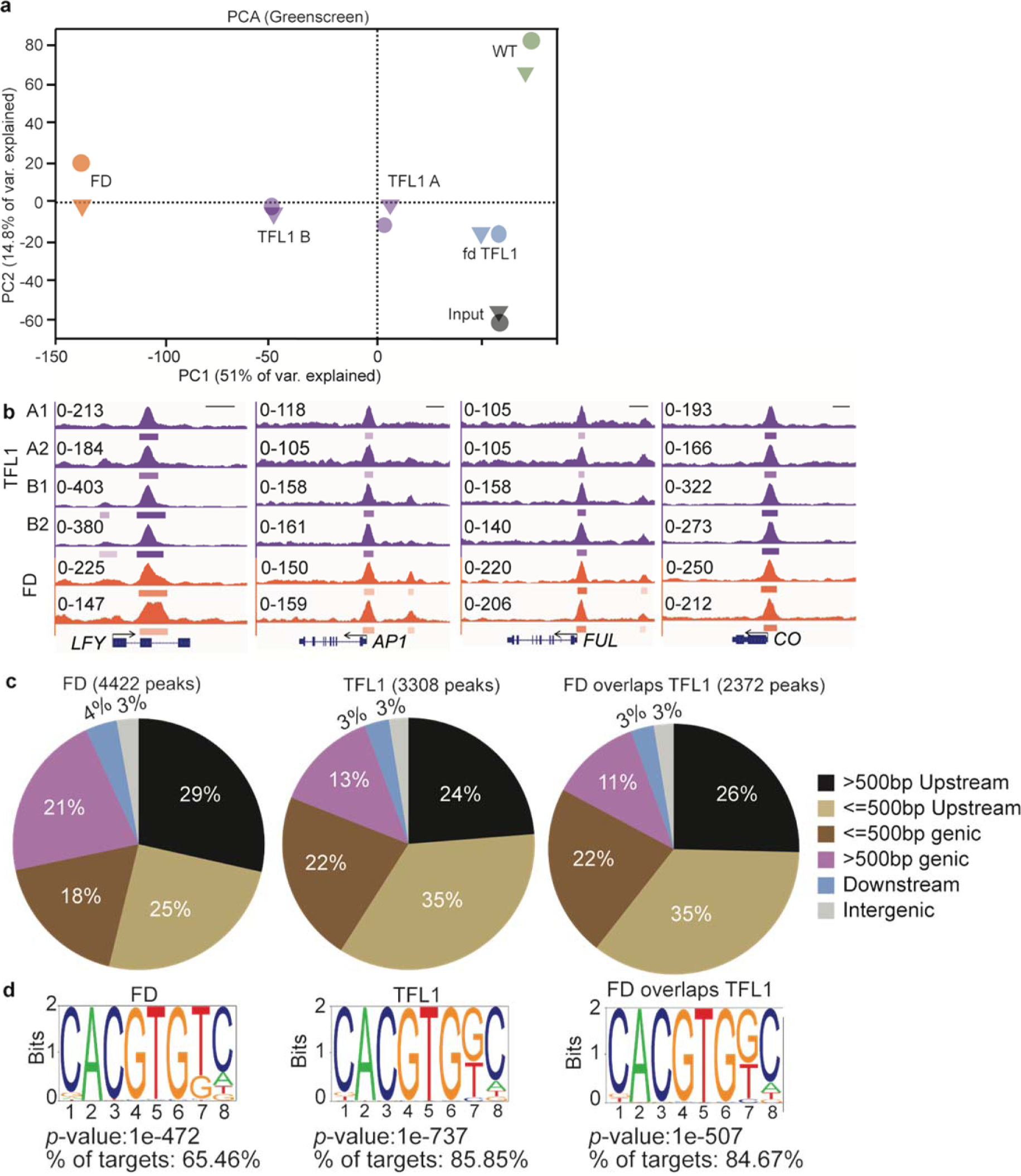
Peak distribution and *cis* motifs for significant TFL1, FD and overlapping ChIP-seq peaks. **a**, Principal component analysis (PCA) of CP10M normalized ChIP-seq reads from anti-GFP ChIP in gTFL1-GFP (TFL1 A and B), *fd-1* gTFL1-GFP (*fd* TFL1), and wild type (WT; negative control), anti-GUS ChIP-seq reads for gFD-GUS (FD), and input. **b**, Browser view of peaks in individual ChIP replicates for TFL1 (A1, A2, B1 and B2) and FD. Significant peaks (summit MACS2 q value < 10^−10^) are marked by horizontal bars, with the color saturation proportional to the -log 10 q value (as for the narrowPeak file format in ENCODE). **c**, ChIP-seq peak summit distribution of FD, TFL1 and the overlapping peaks (TFL1 and FD) in genic and intergenic regions. Intergenic peaks are > 4kb from the closest gene. **d**, *cis* motifs most significantly enriched under TFL1 or FD peaks identified by *de novo* motif analysis (Homer ^43^). Sequence logos of position-specific scoring matrices (PSSMs) (top). Motif enrichment p-values and frequency (bottom).

**Supplementary Figure 3.**
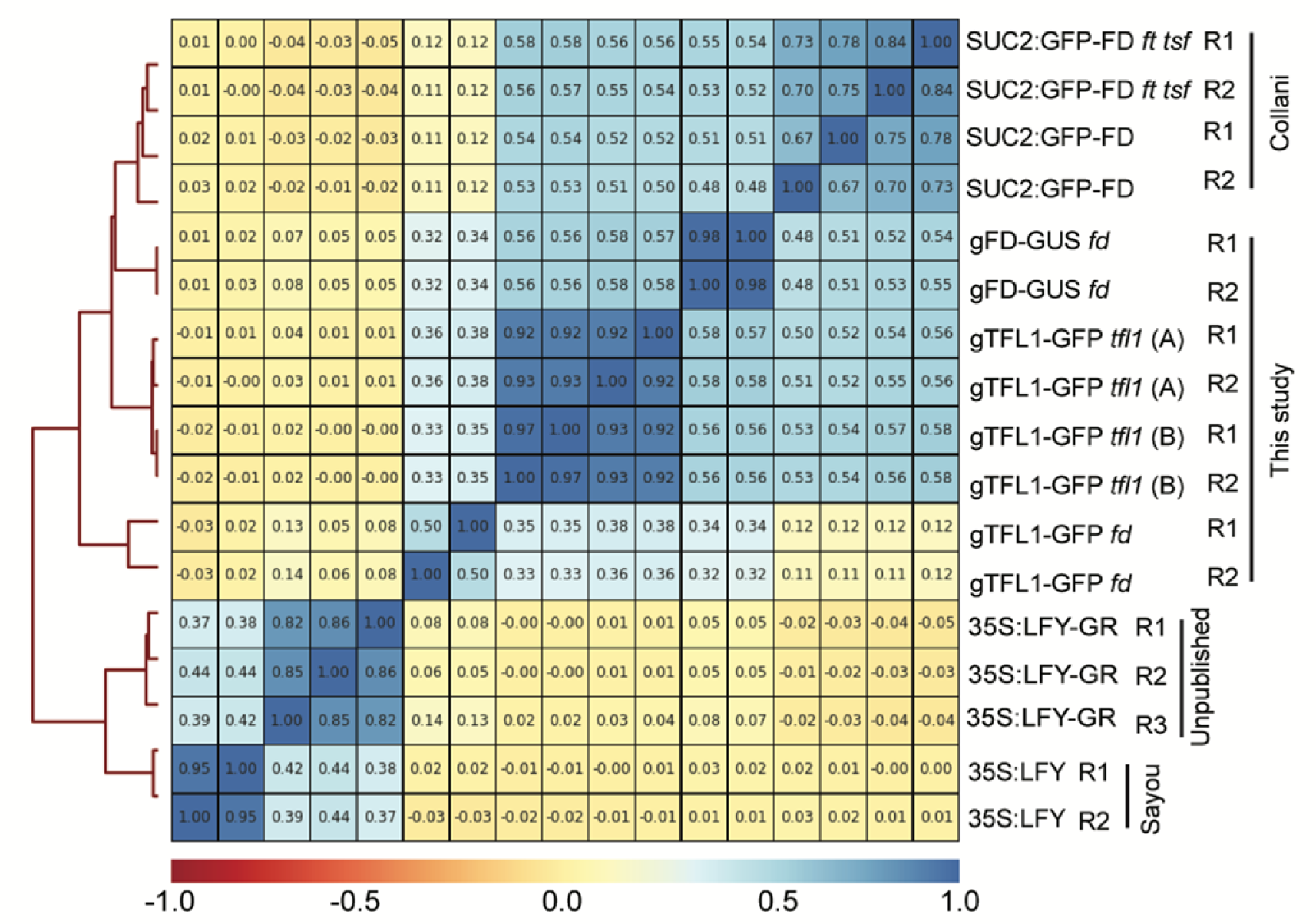
ChIP-seq quality control. Top: Heatmap of Pearson correlation analysis of normalized reads in significant peak regions (MACS2 ≤ q value10^-10^). Bottom: legend for correlation co-efficients. See supplemental methods for details. TFL1 and FD datasets generated in the current study (42-day-old short-day-grown plants; this study) clustered together. They also clustered with a recently published SUC2:GFP-FD ChIP-seq dataset ^48^ (21 day-old short-day grown plants; Collani). An unpublished LFY ChIP-seq experiment from our lab (root explants, 35S:LFY-GR), by contrast, clustered with a published LFY ChIP-seq experiment (inflorescences, 35S:LFY) ^49^, suggesting no strong batch effects.

**Supplementary Figure 4.**
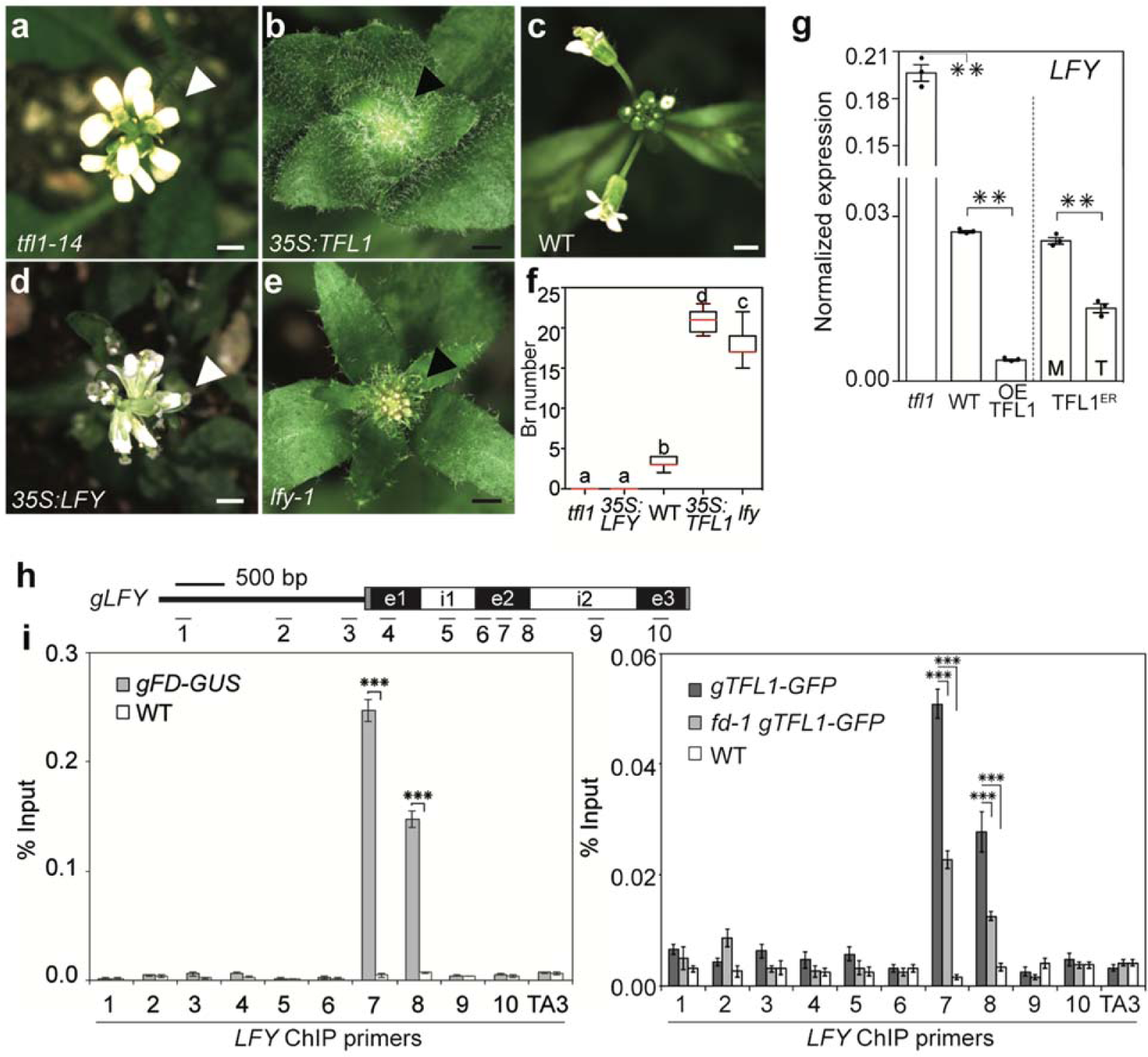
Direct repression of *LFY* by TFL1. **a** - **f**, *TFL1* loss of function phenocopies *LFY* gain-of function and vice versa. Representative pictures of inflorescences (top view) grown in long-day photoperiod (a - e). Quantification of the number of branches (Br) formed (f). Scale bars: 2 mm (white) or 5 mm (black). Box plot: median (red line; n = 12 plants), upper and lower quartiles (box edges), and minima and maxima (whiskers). Letters above boxes indicate significantly different groups; *p*-value < 0.05 based on Kruskal-Wallis test with Dunn’s *post hoc* test. **g**, *LFY* expression in 16-day-old long-day-grown *TFL1* loss- and gain-of function mutants (left) and 4 hours after mock (M) or estradiol (T) induction of TFL1^ER^ (right). Shown are mean ± SEM of three independent experiments (black dots). ** *p*-value < 0.01; unpaired one-tailed *t*-test. OE: 35S:TFL1, Expression was normalized over *UBQ10*. **h**, Diagram of *LFY* locus and ChIP primers used. **i**, Anti-GUS (left) or anti-GFP (right) ChIP of FD binding (left) and TFL1 binding (right), respectively, to the *LFY* locus in 42-day-old short-day-grown shoot apices. TFL1 binding was assayed in the presence and absence of *FD* (*fd-1* null mutant; bottom). The following controls were employed: genomic control: anti-GUS (left) or anti-GFP (right) ChIP of non-transgenic wild type (WT), internal control: occupancy at the TA3 retrotransposon locus. Shown are mean ± SEM of three independent experiments (black dots). *** *p*-value < 0.001; unpaired one-tailed *t*-test.

**Supplementary Figure 5.**
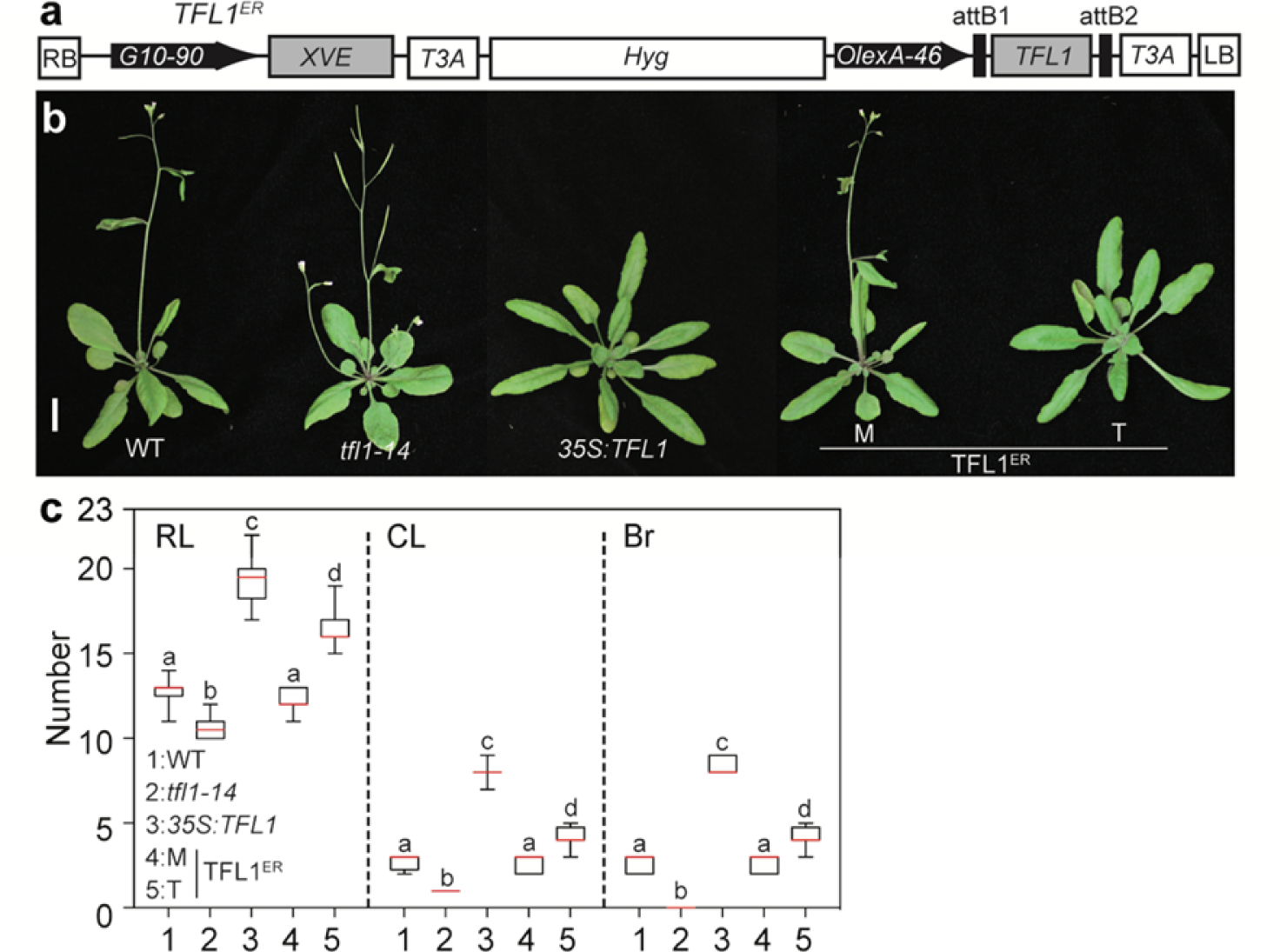
Steroid activatable TFL1 overexpression delays onset of reproductive development and flower formation. **a**, Construction of estradiol inducible TFL1 (TFL1 ^ER^) using pMDC7 (Ref. ^14^). Estradiol application activates the synthetic transcription factor XVE expressed from a constitutive promoter (G10-90). XVE consists of a LexA DNA binding domain (X), a VP16 activation domain (V) and an estradiol receptor hormone binding domain (E). Steroid activated XVE binds to OlexA-46 (eight copies of the *LexA* operator) in front of *TFL1* and activates gene expression. T3A: *rbcsS3A* poly(A) sequence. RB: right border, LB: left border, Hyg: hygromycin B resistance gene. **b**, Phenotypes of long-day-grown plants with decreased (*tfl1-14*), constitutively increased (35S:TFL1) or inducibly increased (TFL1^ER^) TFL1 activity. TFL1^ER^ plants were treated with 10 μmol beta-estradiol (T) or mock (M) solution from day 5 onward every other day until bolting. **c**, Quantification of phenotypes in (b). RL: rosette leaf number, CL: cauline leaf number, Br: branch number on the main inflorescence. Box plot-median (red line; n = 15 plants), upper and lower quartiles (box edges), and minima and maxima (whiskers). Letters above boxes indicate significantly different groups *p*-value < 0.05 based on Kruskal-Wallis test with Dunn’s *post hoc* test.

**Supplementary Figure 6.**
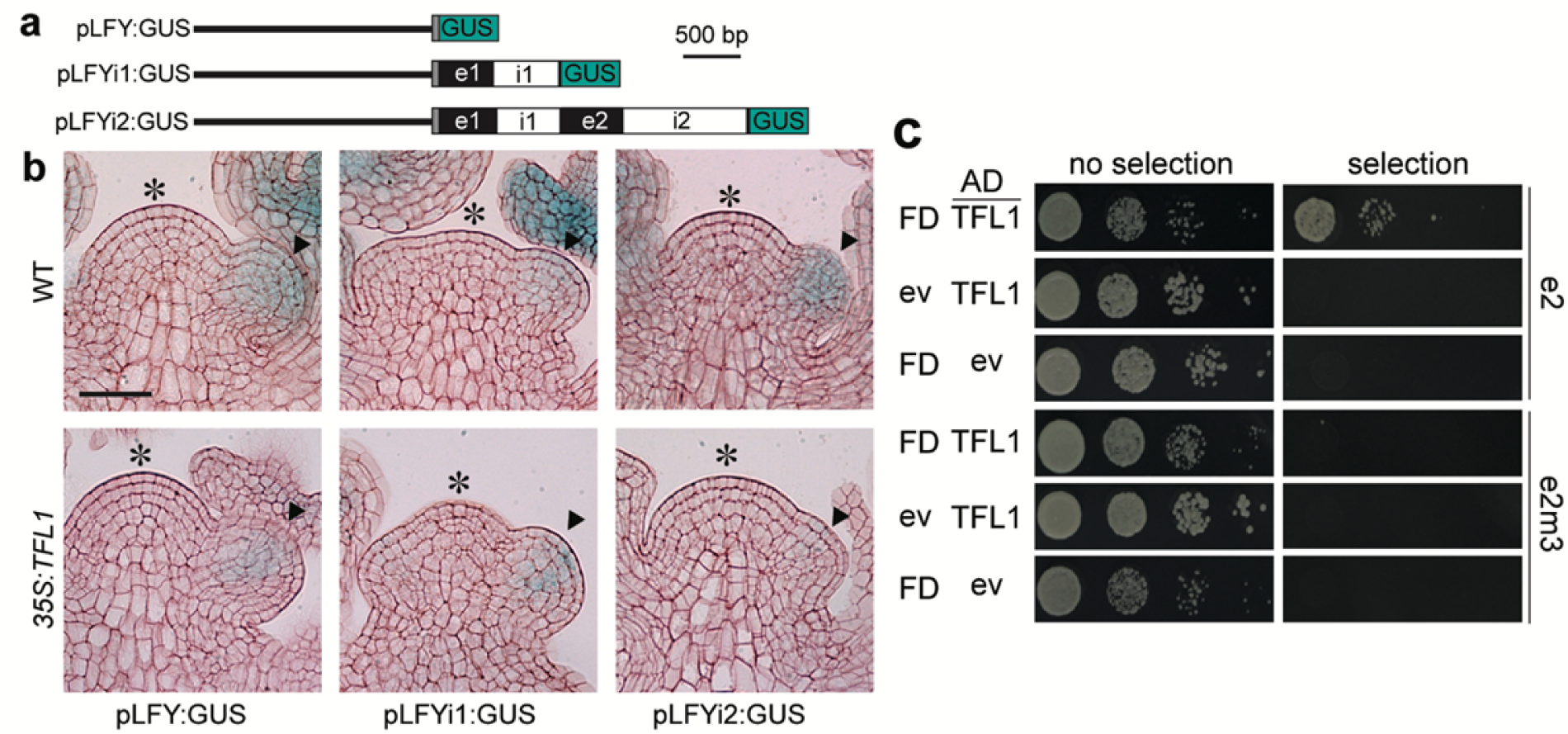
*LFY* locus regions required for repression by and recruitment of TFL1. **a**, LFY-GUS reporter constructs employed. Black line: 2.3 kb upstream intergenic region (*LFY* ‘promoter’) ^16^. Grey box: 5’UTR. Black boxes: exons. White boxes: introns. Blue box: beta-glucuronidase (GUS). **b**, GUS staining in wild-type (WT) and in *35S:TFL1* inflorescences grown in long-day photoperiod. Asterisk indicates the center of the inflorescence shoot apex. Arrowheads point to flower primordia, where *LFY* is expressed. Only the pLFYi2:GUS reporter is silenced by 35S:TFL1. Scale bar: 1 µm. **c**, Test for TFL1/FD binding to wild-type (e2) and bZIP binding site mutated LFY exon 2 (e2m3) in yeast. For details on the *cis* motif mutations see Fig. 2a. LFY exon 2 or e2m3 driving expression of a resistance marker were integrated into the yeast genome. The resulting strains were transformed with plasmids expressing FD and TFL1-AD or control (ev= empty vector) ND TFL1-AD. AD: activation domain. Yeast 14-3-3 proteins mediate FD-florigen interactions in yeast ^60, 61^. Growth was assayed on media without or with selection.

**Supplementary Figure 7.**
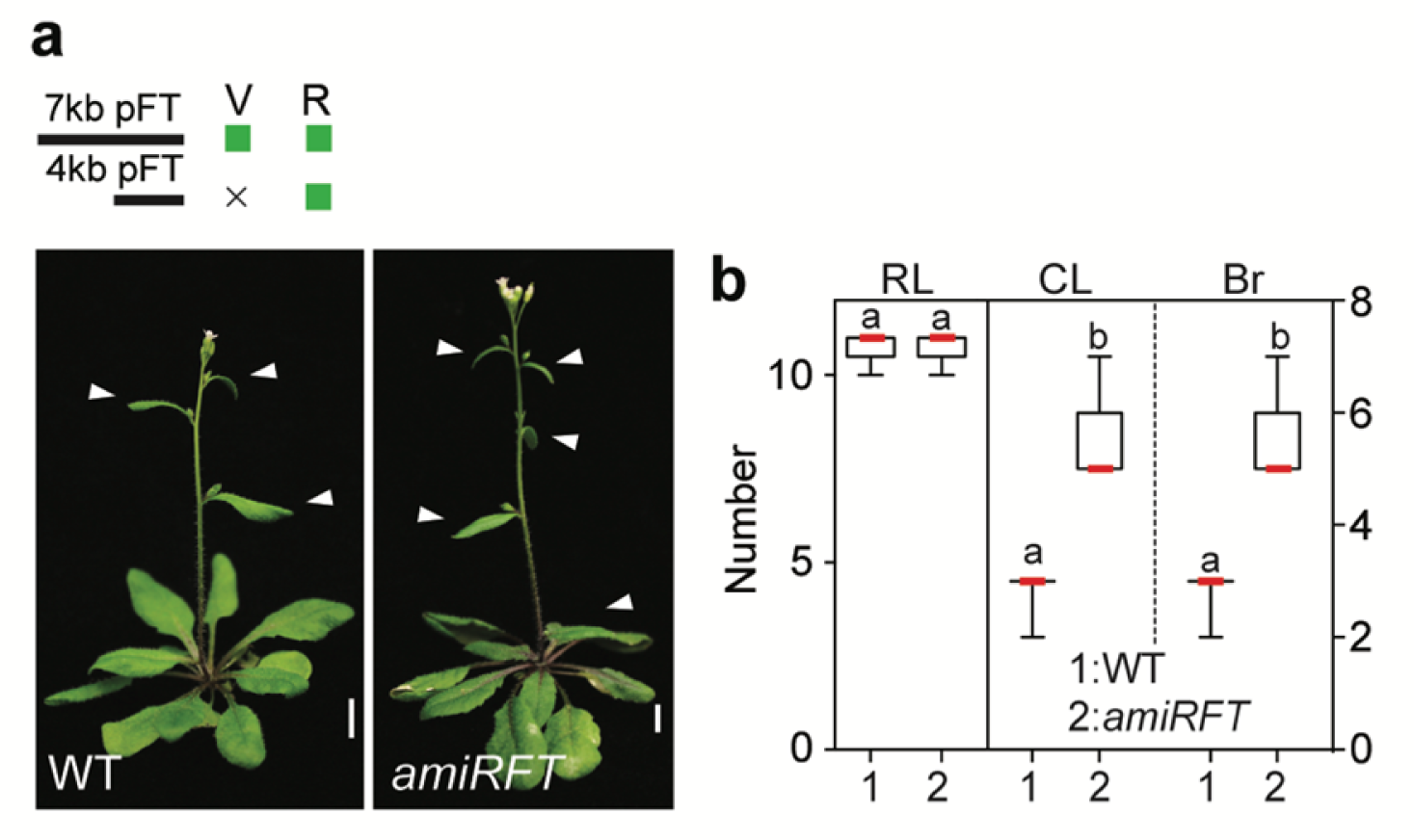
Downregulation of *FT* specifically during reproductive development. **a, b,** Representative images of the phenotypes of *pFT4kb:amiRFT* plants, in which an artificial microRNA specific to *FT* ^22^ was expressed from a minimal *FT* promoter active only during the reproductive phase ^21^. V: vegetative phase, R: reproductive phase (a). Plants were grown in long-day photoperiod. *pFT4kb:amiRFT* plants did not delay onset of reproduction (plants formed the same number of rosette leaves (RL) as the wild type (b). However, the switch to flower formation occurred significantly later (more cauline leaves (CL) and branches (Br) formed) relative to the wild type (b). Quantification, n = 15 plants. Box plot-median (red line), upper and lower quartiles (box edges), and minima and maxima (whiskers). Letters: significantly different groups *p*-value < 0.05 based on Kruskal-Wallis test with Dunn’s *post hoc* test. Scale bars, 1 cm.

**Supplementary Figure 8.**
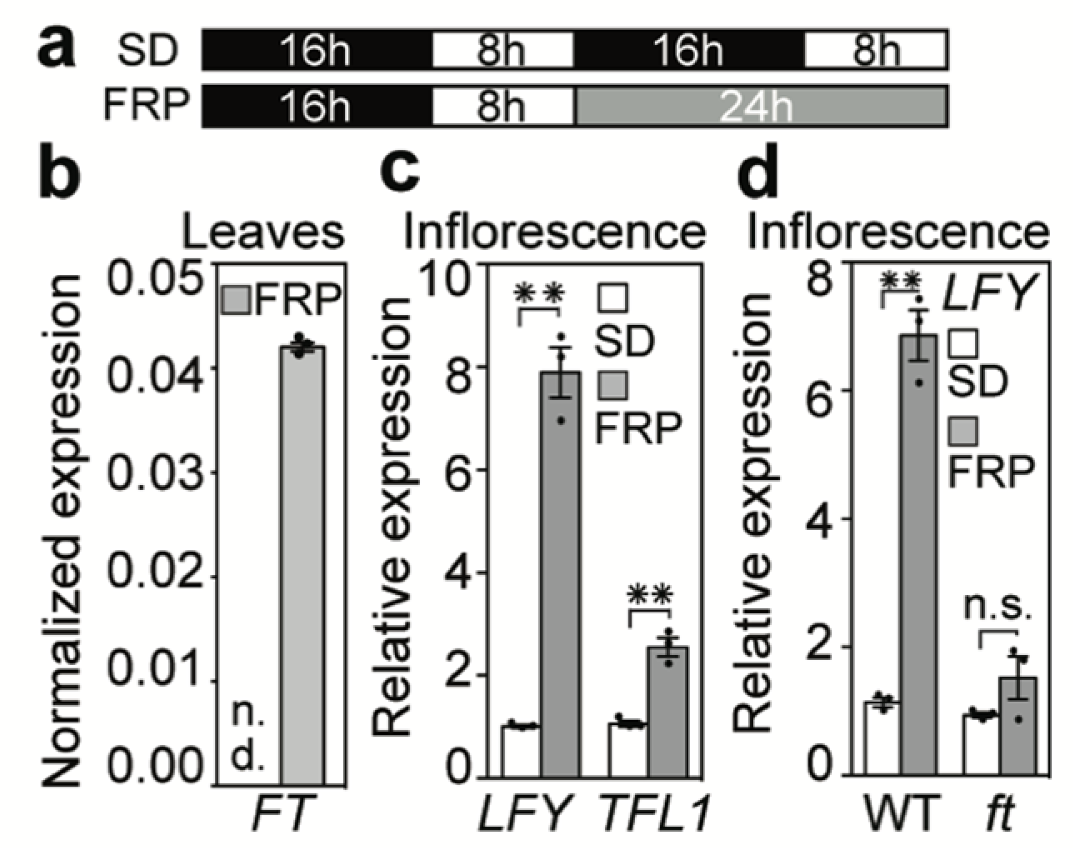
FT promotes *LFY* accumulation. **a**, Diagram for photoperiod triggered upregulation of *FT* in 42-day-old short-day-grown plants. **b - d**, Change in gene expression after far-red light enriched photoinduction (FRP). Gene expression was normalized to that of *UBQ10*. *FT* expression is strongly elevated in fully expanded true leaves (b) which results in FT protein accumulation at the shoot apex ^62^. (c) FRP caused strong upregulation of *LFY* expression in inflorescences. *TFL1* is moderately upregulated consistent with the known increase in *TFL1* levels at the onset of the reproductive phase ^63^. (d) No significant *LFY* upregulation by FRP in the *ft-10* null mutant (see also **Supplementary Fig.12a**). (b - d) Expression is normalized over that of *UBQ10.* (c, d) Normalized gene expression relative to SD grown plants. The normalized values were (mean ± SEM): *LFY*: SD (0.0024±0.0001), FRP (0.0185±0.0012); TFL1: SD (0.0293±0.0016), FRP (0.0703±0.0051) for (c) and *LFY* in SD WT (0.0025±0.0002), *LFY* in FRP WT (0.0152±0.0015); *LFY* in SD *ft* (0.0019±0.0001), *LFY* in FRP *ft* (0.0031±0.0012) for (d). Shown are mean ± SEM of three independent experiments (black dots). * *p*-value < 0.05, ** *p*-value < 0.01; unpaired one-tailed *t*-test in (c, d). n.s.: not significant (*p*-value > 0.05). n.d.: not detectable.

**Supplementary Figure 9.**
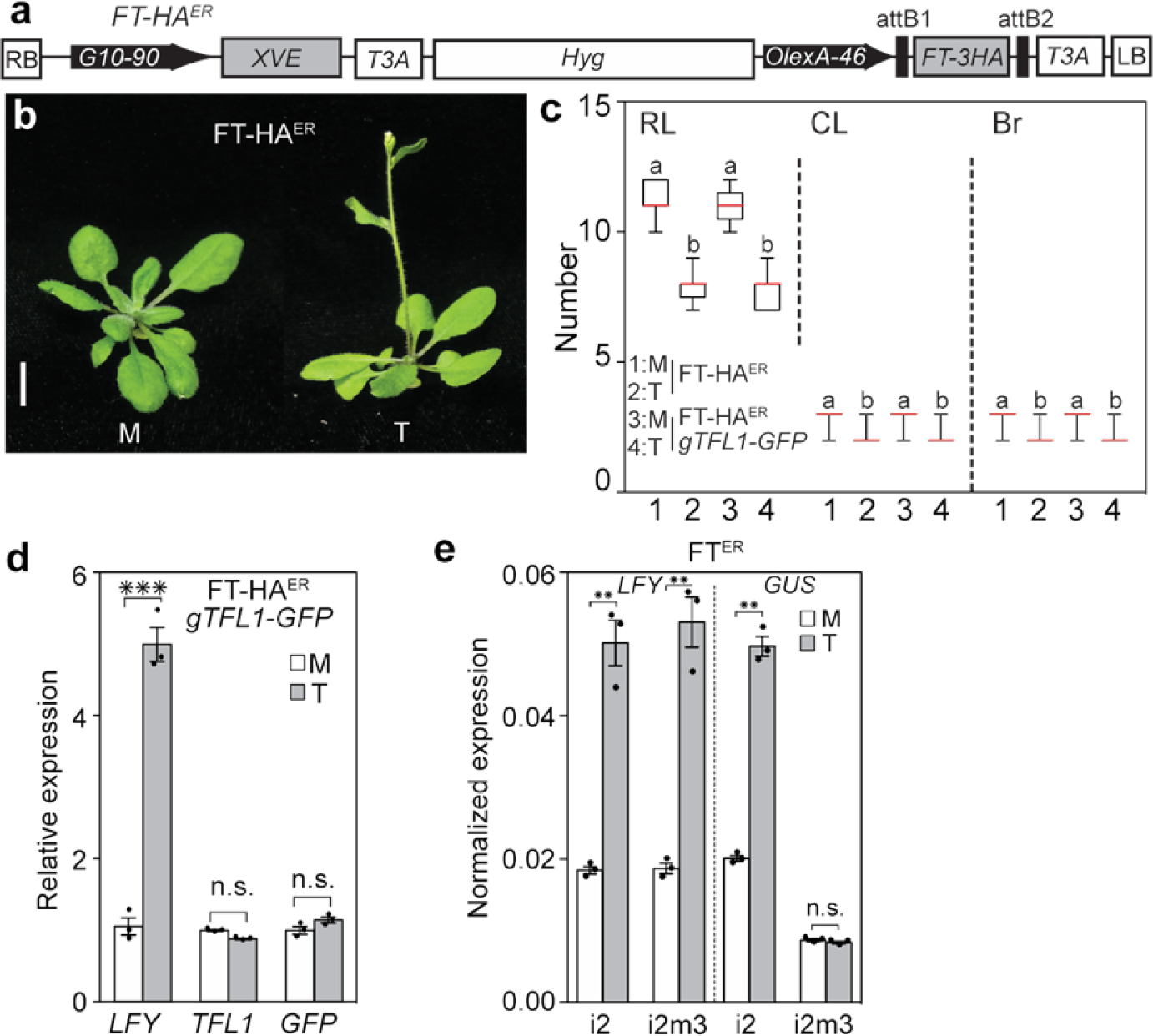
Steroid activated FT causes precocious onset reproductive development. **a**, FT-HA^ER^ construct, see Supplementary Figure 4a for details. **b**, Early flowering phenotypes of plants with inducible increase of FT^ER^. Plants were treated with 10 μmol beta-estradiol (T) or mock (M) solution from day 5 onward every other day until bolting. Scale bars: 1 cm. **c**, Quantification of phenotypes in (b). RL: rosette leaf number, CL: cauline leaf number, Br: branch number. M: mock, T: steroid treatment. Box plot-median (red line; n = 15 plants), upper and lower quartiles (box edges), and minima and maxima (whiskers). Letters above boxes indicate significantly different groups *p*-value < 0.05 based on Kruskal-Wallis test with Dunn’s *post hoc* test. **d**, FT-HA^ER^ activation leads to increased *LFY* accumulation in 12-day-old long-day grown plants, while endogenous *TFL1* or transgene *gTFL1*-*GFP* levels do not change. Gene expression was normalized to *UBQ10* and is displayed relative to the mock (M) condition for better comparison. Normalized values (mean±SEM) were: *LFY* (0.0025±0.0003 (M), 0.0118±0.0006(T)), *TFL1* (0.0150±0.0006 (M), 0.0136±0.0002 (T)), and GFP (0.0059±0.0003 (M), 0.0068 ± 0.0002 (T)). **e**, Effect of FT induction by steroid treatment (4 hrs) in 12-day-old long-day-grown plants on endogenous *LFY* (left) or on a *LFY* reporter (right). Both wild type (pLFYi2:GUS) and bZIP binding site mutated (pLFYi2m3:GUS) reporters were assayed. Expression was normalized over that of *UBQ10.* (d, e) Shown are mean ± SEM of three independent experiments (black dots). *** *p*-value < 0.001; ** *p*-value < 0.01; n.s.: not significant (*p*-value > 0.05) unpaired one-tailed *t*-test.

**Supplementary Figure 10.**
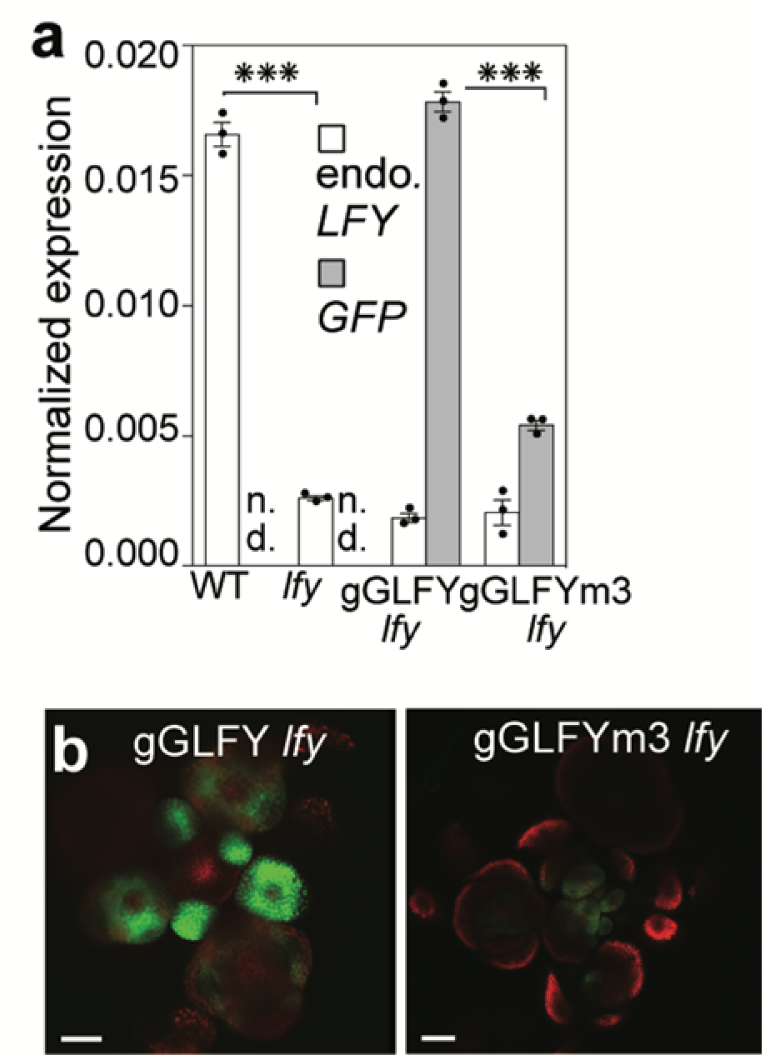
bZIP binding site mutations reduce *LFY* accumulation. a, b, Expression levels of gGLFY *lfy-1* and gGLFYm3 *lfy-1* null mutants assessed by qRT-PCR (a) or confocal imaging (b). (a) Expression of *GFP* (grey) and endogenous *LFY* (white) was normalized over that of *UBQ10.* Shown are mean ± SEM of three independent experiments (black dots). *** *p*-value < 0.001; unpaired one-tailed *t*-test. n.d.: not detectable. (b) The image for gGLFYm3 was taken at higher gain (1.3x) and laser power (2.7x) than that of gGLFY. Scale bars, 2 mm.

**Supplementary Figure 11.**
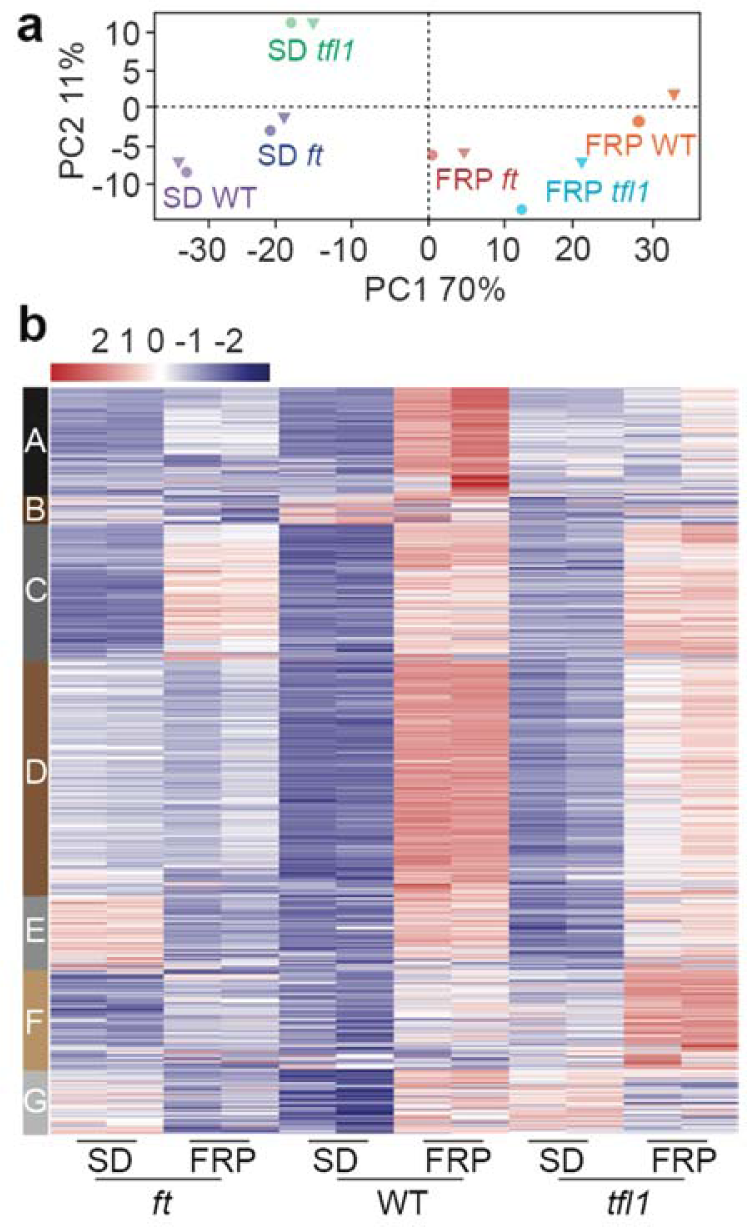
RNA-seq with and without a single far-red enriched photoperiod induction in short-day grown plants. **a**, Principle Component Analysis of normalized reads of dissected inflorescences from *ft-10,* wild-type or *tfl1-1* plants. Plants were grown for 42-days in short-day photoperiod ± photoperiod induction. **b**, K-means clustering of de-repressed genes. DESeq2 normalized expression values are shown for each replicate. Only cluster C lacks FT dependence.

**Supplementary Figure 12.**
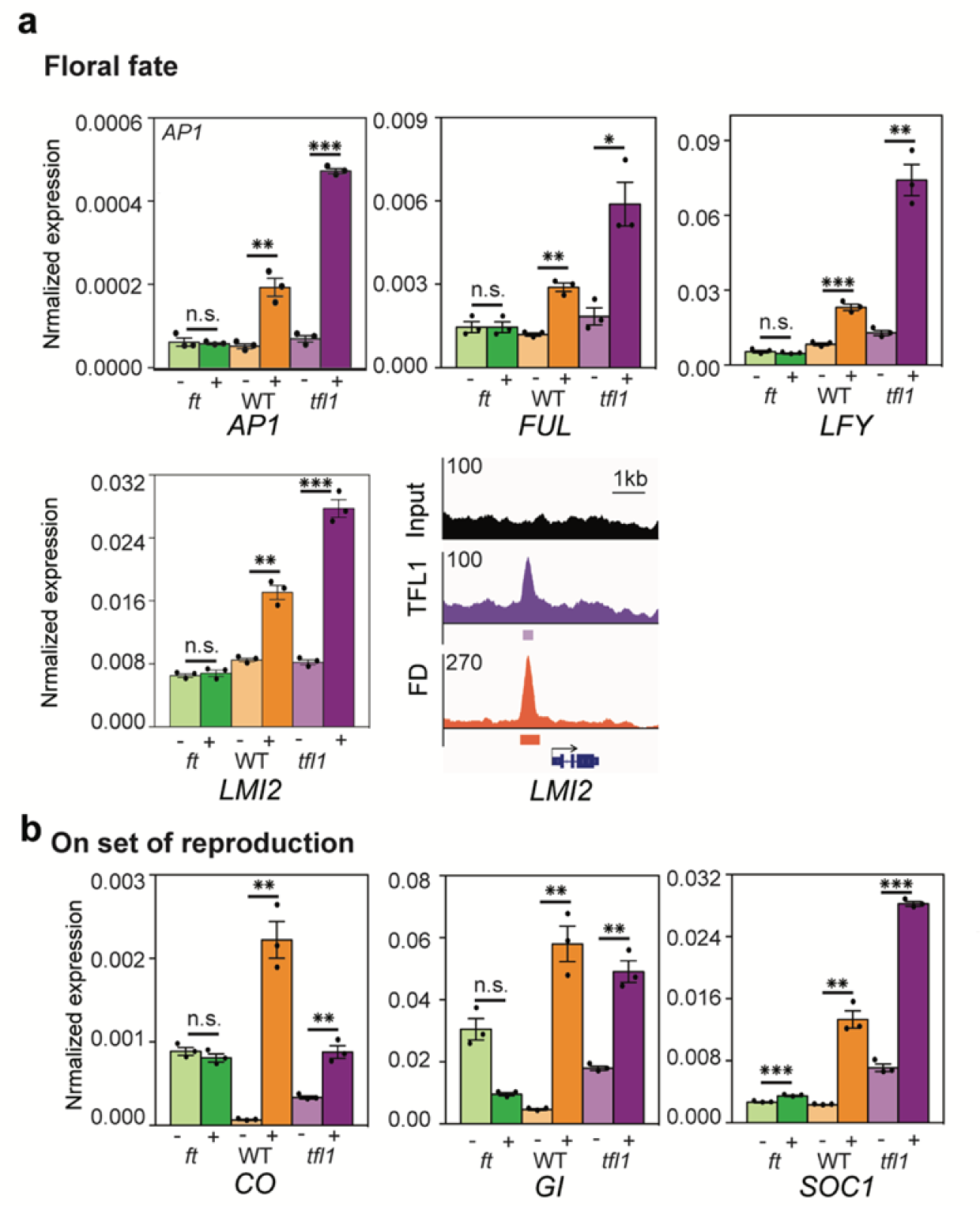
Validation of FT-dependence for TFL1 and FD bound genes involved in onset of reproductive development and flower fate. **a, b,** qRT-PCR of independent biological replicates confirms FT-dependent de-repression of genes directly repressed by the TFL1-FD complex for targets linked to onset of flower fate (a) or onset of reproduction (b). SOC1 alone is significantly de-repressed upon FRP treatment in *ft* mutants. Expression is normalized over that of *UBQ10.* Shown are mean ± SEM of three independent experiments (black dots). * *p*-value < 0.05, ** *p*-value < 0.01, *** *p*-value < 0.001; unpaired one-tailed *t*-test in. n.s.: not significant (*p*-value > 0.05). Browser screenshot of LMI2, which promotes onset of flower formation together with LFY ^64^. Significant peaks (MACS2 summit q value ≤ 10^-10^) are marked by horizontal bars, with the color saturation proportional to the -log 10 q value (as for the narrowPeak file format in ENCODE).

**Supplementary Figure 13.**
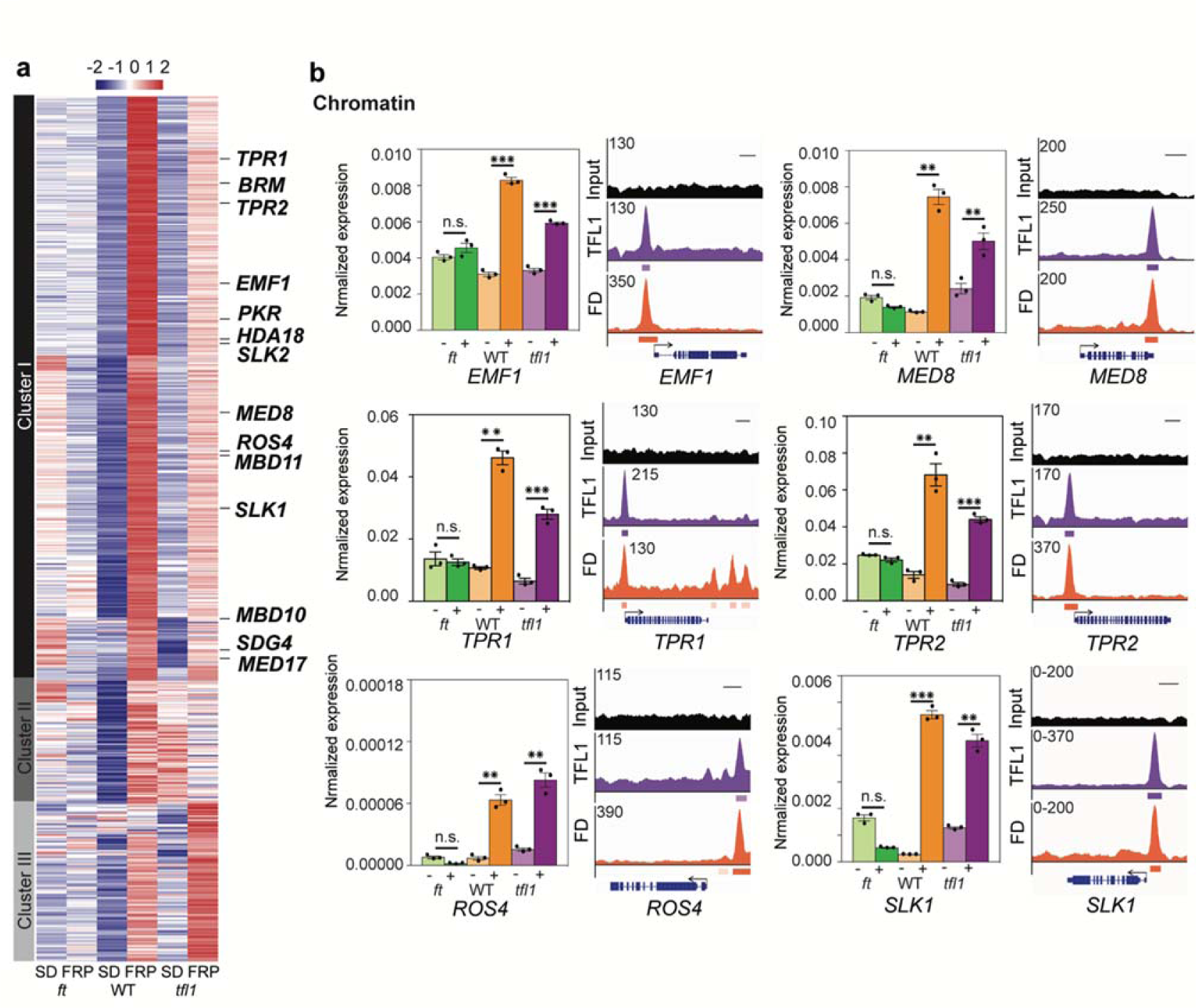
Chromatin regulators bound by TFL1 and FD and de-repressed by FRP. **a**, Chromatin regulators and transcriptional co-regulators targets of TFL-FD complex belong to k-means cluster I. **b,** Right: qRT-PCR of independent biological replicates confirms FT-dependent de-repression of gene directly repressed by the TFL1-FD complex. Expression is normalized over that of *UBQ10.* Shown are mean ± SEM of three independent experiments (black dots). * *p*-value < 0.05, ** *p*-value < 0.01, *** *p*-value < 0.001; unpaired one-tailed *t*-test in. n.s.: not significant (*p*-value > 0.05). Left: Browser screenshots for input, TFL1 and FD ChIP-seq. Significant peaks (MACS2 summit q val ≤ 10^-10^) are marked by horizontal bars, with the color saturation proportional to the -log 10 q value (as for the narrowPeak file format in ENCODE). Direct TFL1-FD targets include mediator complex subunit (*MED*), which links to Pol II transcription ^65^, co-repressor complexes linked to histone de-acetylation such as SEUSS-LIKE (*SLK*) and TOPLESS-RELATED (*TPR*) ^66^ and HISTONE DEACETYLASE 18 (*HDA18*). BRAHMA (*BRM*) is a SWI/SNF chromatin remodeler ^66^, REPRESSOR OF SILENCING 4 (*ROS4*) a histone acetyltransferase ^67^, EMBRYONIC FLOWER 1 (*EMF1*) and PICKLE RELATED 1 (*PKR*) are linked to Polycomb repression ^68^, ASH1-RELATED 3/SET DOMAIN GROUP 4 (*AHR3/SDG4*) is a histone methyltransferase ^69^, METHYL-CPG-BINDING DOMAIN 10 and 11 (*MBD10/11*) are putative methyl-DNA binding proteins ^70^.

**Supplementary Figure 14.**
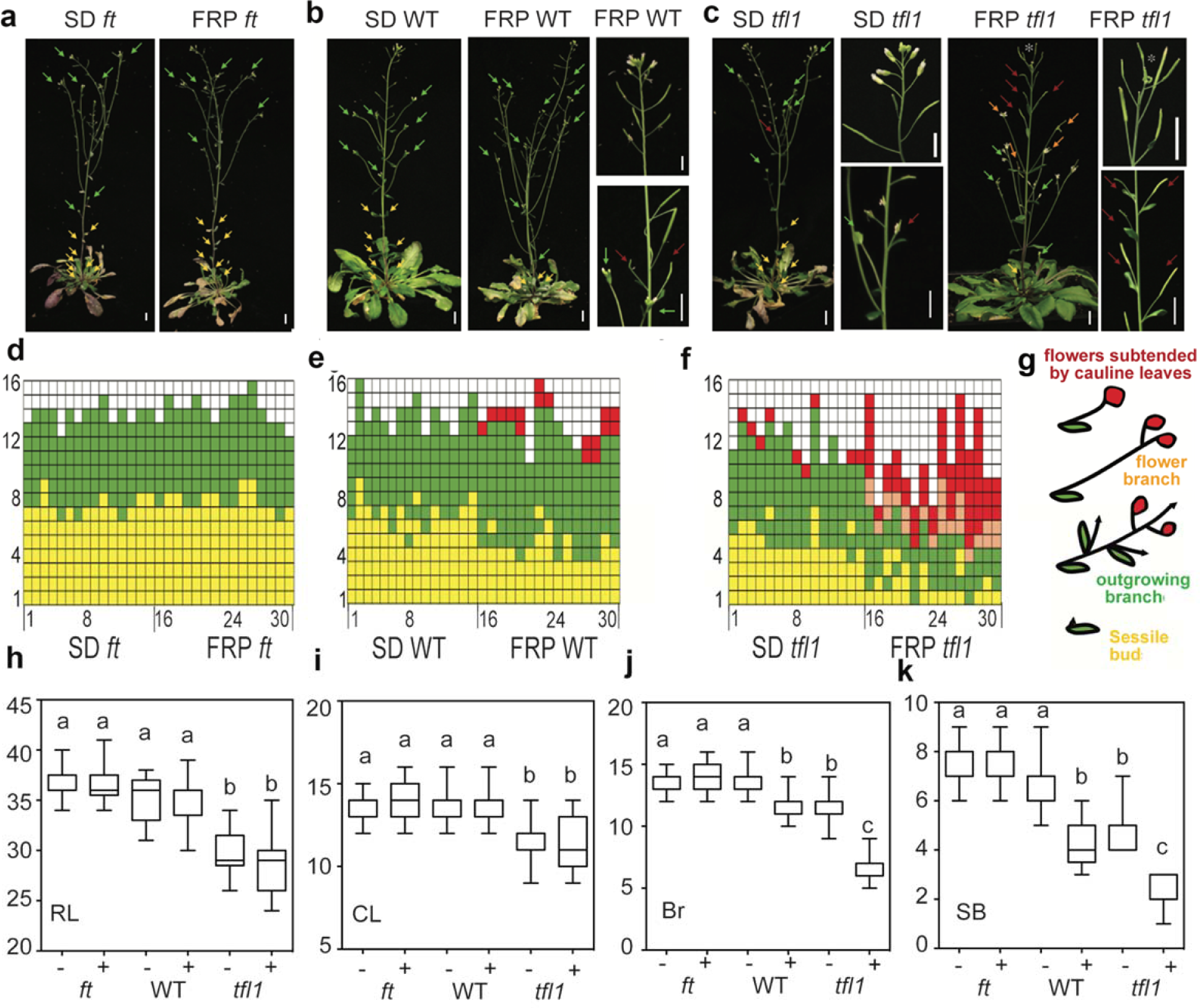
Effect of a single far-red light enriched photoperiod (FRP) on *ft-10* mutants, wild type and *tfl1-1*. **a - c**, Representative plant images. Yellow arrows: sessile buds. Green arrows: outgrowing branches. Orange arrows: flower branches (a single or two flowers borne at the end of a branch-like petiole subtended by a cauline leaf). Red arrows: flowers subtended by cauline leaves. Scoring was terminated when ‘regular’ flowers not subtended by cauline leave formed. Asterisk (*) indicates a terminal flower. Scale bar = 1 cm. **d - f**, Identity and fate of primordia formed on the inflorescence from the bottom (node 1) to the top (node 16) Color coding is as in (a-c): Yellow: sessile buds. Green: outgrowing branches. Orange: flower branches. Red: flowers subtended by cauline leaves. **g,** Schematic of the types of structures formed and color key for (a - f). **h** - **k**, Phenotype quantification. Box plot-median (red line; n = 15 plants), upper and lower quartiles (box edges), and minima and maxima (whiskers). RL, rosette leaves (h), CL: cauline leaves (i), Br, branches (j), SB: sessile buds (k). -, no FRP treatment; +, a single FRP treatment. Letters above boxes indicate significantly different groups (*p*-value < 0.05) based on Kruskal-Wallis test with Dunn’s *post hoc* test. Compared to untreated plants, a single FRP triggered formation of significantly fewer branches in *tfl1* mutants and in the wild type, indicating a switch to floral fate. As the number of cauline leaves, which subtend branches but not flowers in *Arabidopsis*, did not changed, we propose that axillary branch meristems switched to flower fate (as previously proposed by Ref ^71^). In addition, FRP triggered significant branch outgrowth in *tfl1* mutants and the wild type relative to untreated plants (fewer sessile buds formed). Onset of the reproductive phase was unchanged in these plants. This is expected as FRP was applied at day 42, after the plants had terminated vegetative development. No significant effect of FRP treatment was detectable in the *ft* mutant.

**Supplementary Table 1.** Gene Ontology terms enriched in genes with TFL1 and FD binding peaks.

| Description | FDR |
| --- | --- |
| response to abiotic stimulus | 4.50E-35 |
| response to stimulus | 1.30E-30 |
| response to endogenous stimulus | 1.40E-28 |
| regulation of gene expression | 5.20E-22 |
| developmental process | 2.60E-14 |
| anatomical structure development | 3.20E-13 |
| response to stress | 1.90E-12 |
| photosynthesis | 1.40E-10 |
| post-embryonic development | 3.50E-10 |
| cell communication | 9.90E-10 |
| signal transduction | 2.70E-09 |
| cellular biosynthetic process | 1.30E-08 |
| reproductive structure development | 4.70E-08 |
| generation of precursor metabolites and energy | 1.50E-06 |
| flower development | 5.90E-06 |

Significant Peaks (MACS2 q value $\leq 10^{-10}$ ) were annotated to genes as described in the Materials and Methods. GO slim was implemented in AgriGO v2.0 (Ref. <sup>46</sup>) and GO terms associated with Yekutieli (FDR under dependency) significance level $< 10^{-5}$ were identified.

**Supplementary Table 2.**
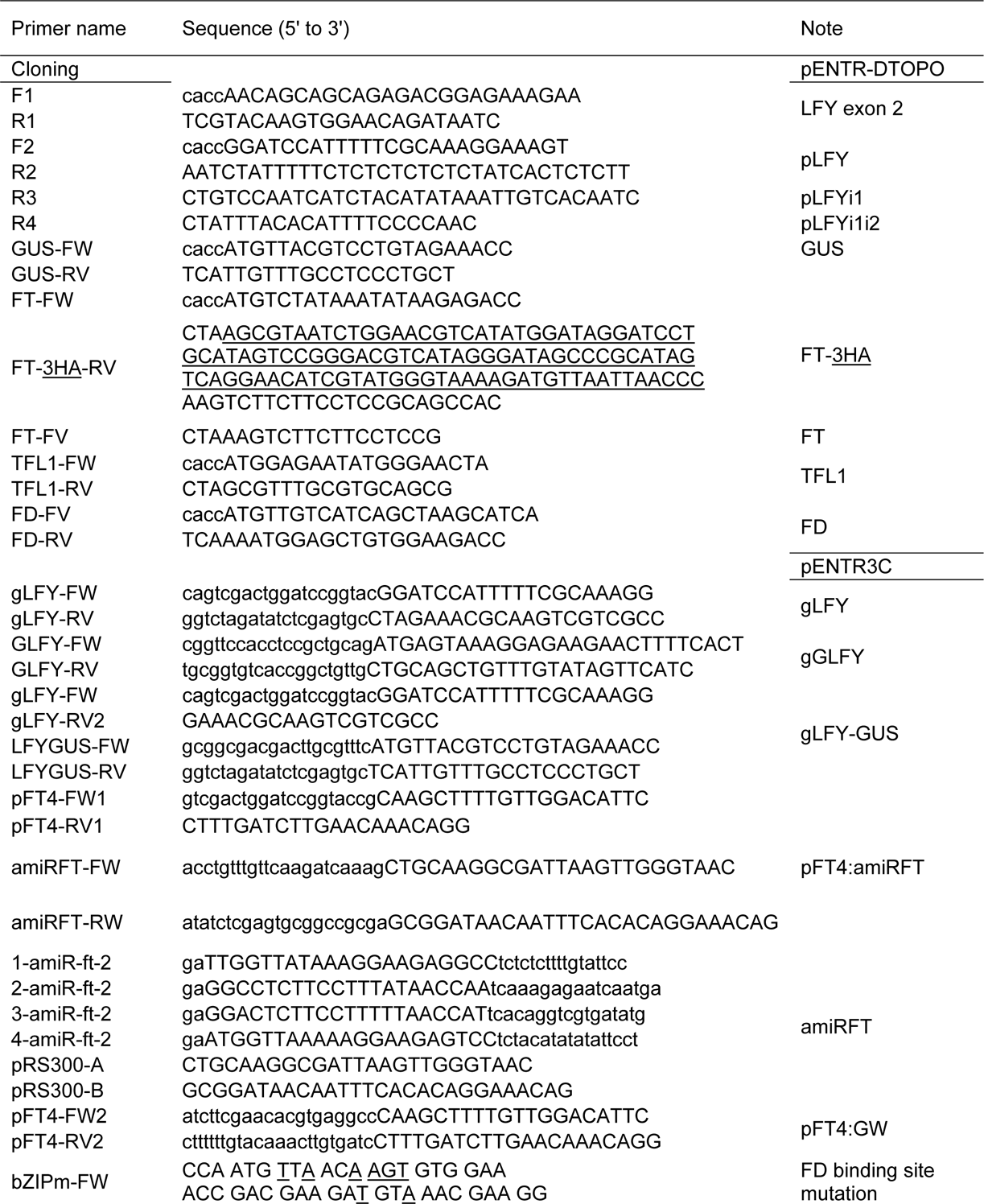
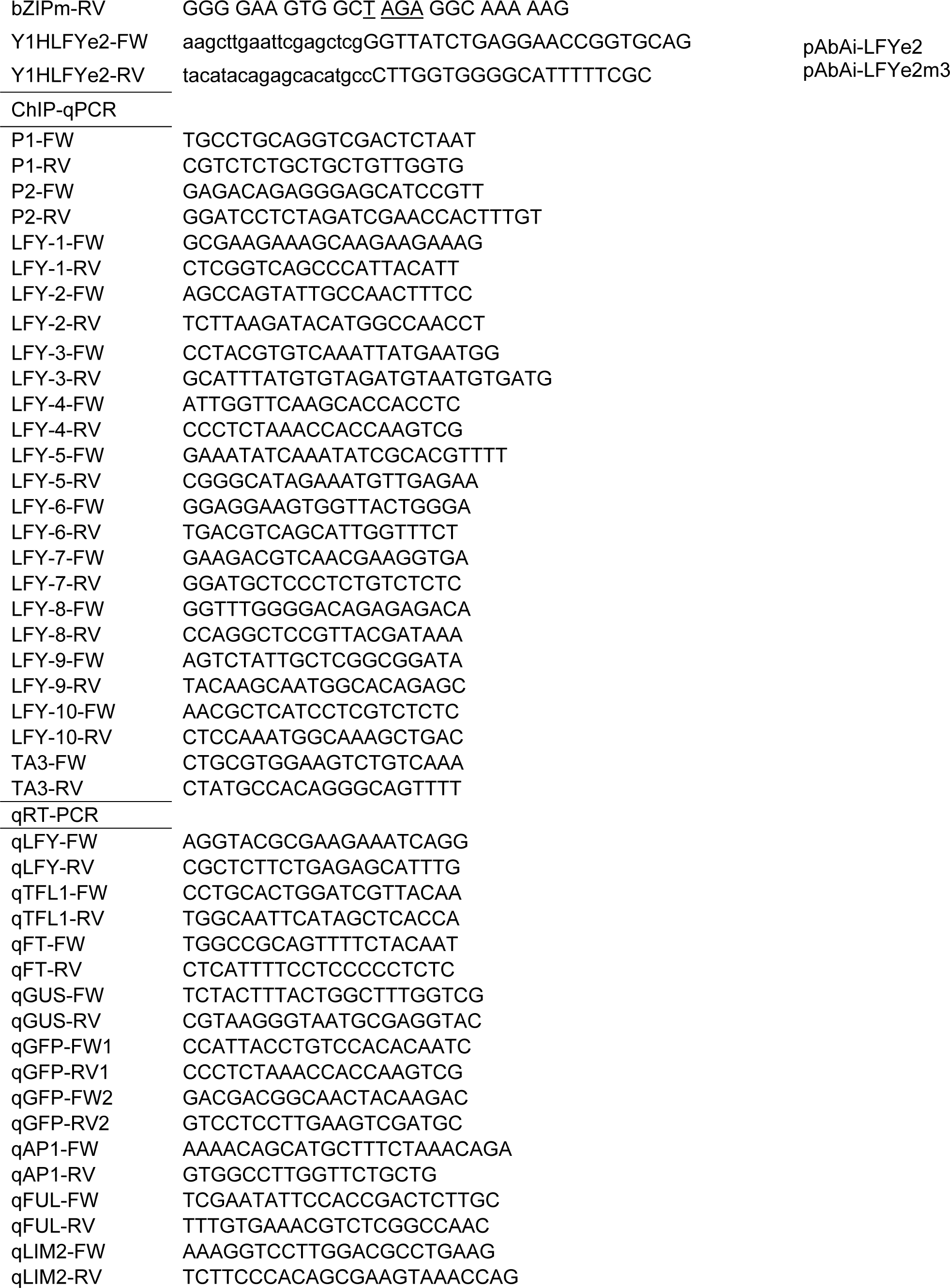
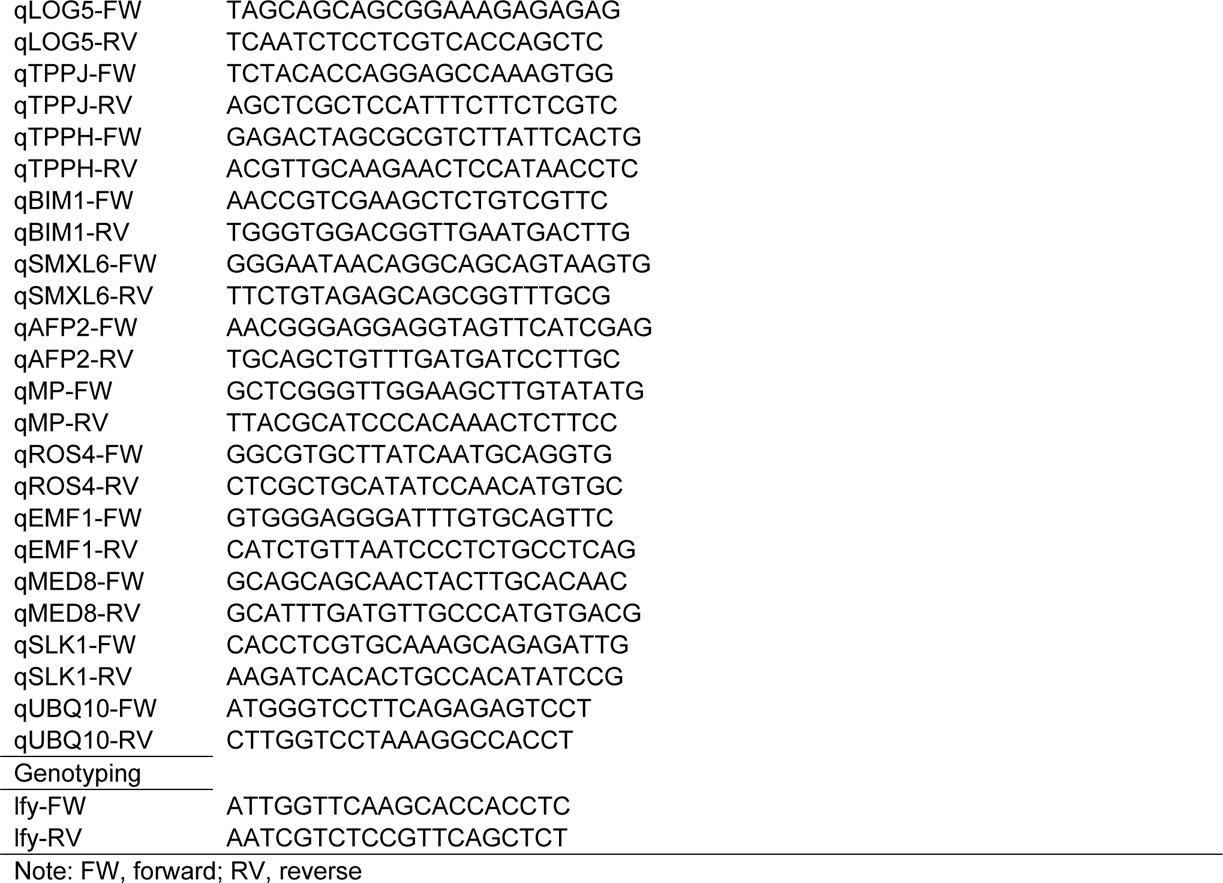
Primers used

**Supplementary Table 3.** Summary of plant lines generated.

| Construct | Transformed into | Crossed to |
| --- | --- | --- |
| *gTFL1-GFP* | *tfl1-1* |  |
| *TFL1<sup>ER</sup>* | wild type |  |
| *FT-HA<sup>ER</sup>* | wild type | *pLFYi2:GUS*<br>and *pLFYi2m3:GUS* |
| *FT-HA<sup>ER</sup>* | *gTFL1-GFP tfl1-1* |  |
| *pLFY:GUS* | wild type |  |
| *pLFYi1:GUS* | wild type |  |
| *pLFYi2:GUS* | wild type |  |
| *pLFYi2m3:GUS* | wild type |  |
| *e2* | wild type<br>*gTFL1-GFP tfl1-1*<br>*FT-HA<sup>ER</sup>* |  |
| *e2m3* | wild type<br>*gTFL1-GFP tfl1-1*<br>*FT-HA<sup>ER</sup>* |  |
| *pFT4kb:amiRFT* | wild type |  |
| *gGLFY* | *lfy-1/+<sup>\*</sup>* |  |
| *gGLFYm3* | *lfy-1/+<sup>\*</sup>* |  |
| *gLFY:GUS* | wild type |  |
| *gLFYm3:GUS* | wild type |  |

\* Heterozygous *lfy-1 -/+* seed stock was generated by crossing *lfy-1* to line C2723 (Ref.
<sup>72</sup>), which contains two fluorescent markers flanking the *lfy-1* locus (*NAP:eGFP*
*NAP:dsRED*, C2723). Homozygous *lfy-1* plants were identified in the T2 generation by
fluorescence marker selection and confirmed by genotyping.

